# Phasing millions of samples achieves near perfect accuracy, enabling parent-of-origin classification of variants

**DOI:** 10.1101/2024.05.06.592816

**Authors:** Cole M. Williams, Jared O’Connell, William A. Freyman, 23andMe Research Team, Christopher R. Gignoux, Sohini Ramachandran, Amy L. Williams

**Affiliations:** Center for Computational Molecular Biology, Brown University, Providence RI; Department of Ecology, Evolution, and Organismal Biology, Brown University, Providence RI; 23andMe, Inc., Sunnyvale CA; Colorado Center for Personalized Medicine, University of Colorado Anschutz, Aurora CO; Data Science Institute, Brown University, Providence RI

## Abstract

Haplotype phasing, the process of determining which genetic variants are physically located on the same chromosome, is crucial for various genetic analyses. In this study, we first benchmark SHAPEIT and Beagle, two state-of-the-art phasing methods, on two large datasets: > 8 million diverse, research-consented 23andMe, Inc. customers and the UK Biobank (UKB). We find that both perform exceptionally well. Beagle’s median switch error rate (SER) (after excluding single SNP switches) in white British trios from UKB is 0.026% compared to 0.00% for European ancestry 23andMe research participants; 55.6% of European ancestry 23andMe research participants have zero non-single SNP switches, compared to 42.4% of white British trios. South Asian ancestry 23andMe research participants have the highest median SER amongst the 23andMe populations, but it is still remarkably low at 0.46%. We also investigate the relationship between identity-by-descent (IBD) and SER, finding that switch errors tend to occur in regions of little or no IBD segment coverage.

SHAPEIT and Beagle excel at ‘intra-chromosomal’ phasing, but lack the ability to phase across chromosomes, motivating us to develop an inter-chromosomal phasing method, called HAPTIC (**HAP**lotype **TI**ling and **C**lustering), that assigns paternal and maternal variants discretely genome-wide. Our approach uses identity-by-descent (IBD) segments to phase blocks of variants on different chromosomes. HAPTIC represents the segments a focal individual shares with their relatives as nodes in a signed graph and performs bipartite clustering on the signed graph using spectral clustering. We test HAPTIC on 1022 UKB trios, yielding a median phase error of 0.08% in regions covered by IBD segments (33.5% of sites). We also ran HAPTIC in the 23andMe database and found a median phase error rate (the rate of mismatching alleles between the inferred and true phase) of 0.92% in Europeans (93.8% of sites) and 0.09% in admixed Africans (92.7% of sites). HAPTIC’s precision depends heavily on data from relatives, so will increase as datasets grow larger and more diverse. HAPTIC enables analyses that require the parent-of-origin of variants, such as association studies and ancestry inference of untyped parents.

## INTRODUCTION

Haplotype phasing is the process of determining which genetic variants are located on the same physical chromosome, enabling genotype imputation, local ancestry inference, and identity-by-descent (IBD) calling [28, 27, 13, 35]. It has also been used to study compound heteroyzgosity [16], to boost power in copy-number variant calling [14], and for parent-of-origin effect genome-wide association (GWA) studies [10]. State-of-the-art tools such as SHAPEIT [19] and Beagle [4] can now phase hundreds of thousands of samples across entire chromosomes. However, their performance has primarily been assessed using trio children from the UK Biobank, most of which are of white British ancestry (Beagle was further assessed with two admixed African-ancestry groups) [4]. SHAPEIT and Beagle employ the Li & Stephens copying-with-recombination model [25] implemented through hidden Markov model (HMM) methods to achieve phasing with remarkably high accuracy. Recent advances have further improved algorithmic speed, memory efficiency, and the ability to phase rare variants [19, 4]. Currently, Beagle and SHAPEIT offer the most cost-effective approaches for high-accuracy phasing, although emerging long-read molecular sequencing technologies hold promise for further improvements [1, 8].

However, current phasing methods have an important limitation: they phase each chromosome independently, despite the fact that parents transmit one copy of each chromosome to their children. We use the term ‘intra-chromosomal phasing’ for the analyses SHAPEIT and Beagle employ and ‘inter-chromosomal phasing’ as the process of determining the sets of variants (genome-wide) that an individual co-inherited *across chromosomes*. There has been recent interest in inter-chromosomal phasing as Hofmeister et al. (2021) [20] and Noto & Ruiz (2022) [29] both developed inter-chromosomal phasing algorithms that use genetic relatives and identity-by-descent (IBD). Hofmeister et al. (2021) [20] further demonstrated the utility of inter-chromosomal phasing by performing parent-of-origin effect GWA studies across a range of complex traits. Both these inter-chromosomal phasing methods employ the paradigm described by Kong et al. (2008) [23]: if the parent-of-origin of IBD segments is known, the IBD segments can be used for phasing (i.e., the relative’s genotype data in the region of IBD can be used to perform Mendelian phasing). A key difference between Kong et al. (2008) [23] and recent methods [20, 29] is in the task of assigning relatives (and their IBD segments) parent-of-origin labels. While Kong et al. (2008) [23] make the assignments according to a known genealogy, the recent methods make the assignments without requiring genealogical information.

In this paper, we first benchmark Beagle and SHAPEIT’s performance phasing a subset of trio children against a reference panel of 8.6M research-consented 23andMe, Inc. customers. This panel is 17x larger than the UK Biobank and significantly more diverse. We assess phasing performance in about 6,000 23andMe trio children, which represent six broadly-defined ancestry groups: admixed African, European, Ashkenazi Jewish, South Asian, East Asian, and admixed American. An overall goal in this analysis was to understand the performance gains realized in a large dataset, as well as understand the performance of these methods in non-European ancestry samples. We perform further analyses using 594 trio children from the 1000 Genomes Project [5] and the UK Biobank as a reference panel to characterize best practices for phasing diverse samples while using a reference panel that is primarily European-descent.

Next, we outline and assess our new method for inter-chromosomal phasing, HAPTIC (**HAP**lotype **TI**ling and **C**lustering). Our method takes as input identity-by-descent (IBD) segments computed from pre-phased genotype data (here, the output from SHAPEIT or Beagle). HAPTIC clusters the IBD segments into two sets intended to correspond to maternally-inherited and paternally-inherited IBD segments. We note that in practice determining maternal and paternal labels for the clusters is non-trivial (except in males) without additional genealogical information, but HAPTIC can resolve IBD segments inherited from one “side of the family” versus the other with this input. The output of HAPTIC lists the inferred co-inherited haplotype segments, which may be on opposite haplotypes in the pre-phased data due to switch errors.

Our approach is similar to [29], with a few key differences. Firstly, our method does not require genotype or sequence data and can be run using only IBD segments, avoiding the computational overhead of loading the sequence data. Secondly, [29] use a greedy algorithm to cluster IBD segments, whereas we use spectral clustering on a signed-graph for clustering, as first demonstrated in [24]. Our spectral clustering approach has advantages over a greedy algorithm: the only parameters of the method are the edge weight criteria (which are interpretable summaries of IBD sharing) and the smallest eigenvalue from the eigen-decomposition can be an indicator of performance. We validate HAPTIC using the same ∼6,000 trio children from the 23andMe cohort from six broad population groups and ∼1,000 trio children from the UK Biobank. HAPTIC can be applied to parent-of-origin effect GWA studies (see also [19]), but HAPTIC enables other analyses and inferences that use parental haplotype information.

## METHODS

In our intra-chromosomal phasing analysis, we use three datasets: 8.6M research-consented 23andMe customers, the UK Biobank (487k individuals), and the 1000 Genomes Project (3.2k individuals). For the inter-chromosomal phasing analysis, we used only the UK Biobank and 23andMe samples as 1000 Genomes Project samples do not have the sufficient IBD required for HAPTIC. In all analyses, we evaluated accuracy with trio children.

### 23andMe dataset

The 8.6M 23andMe customers include research-consented individuals genotyped on our most up-to-date SNP array (our ‘v5’ array, which is based on the Illumina Global Screening Array and includes ∼50,000 custom variants). To perform variant-level quality control, we excluded a variant if it lacked GRCh38 coordinates, had a minor allele count (MAC) < 5, or had a missingness rate ≥ 0.1. We further excluded variants if they were not polymorphic in an internal set of 4,893 whole genome sequenced (WGS) research participants or if their genotype calls (encoded as dosage values of 0, 1, or 2) were not well-correlated (Pearson *r <* 0.9) with the corresponding WGS calls. However, we retained these former variants if their allele frequencies and those of the gnomAD non-Finnish European samples [7] were roughly concordant. Specifically, we required their *z*-score (calculated as 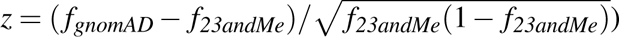 to be in the empirical [0.001, 0.999] quantiles of the normal distribution. The resulting dataset included 608,965 variants across the autosomes and the X-chromosome. Following these variant filters, we excluded individuals whose ratio of the number of heterozygous to the number of homozygous variants is more than 10 standard deviations (SD) above the median (using the median absolute deviation to estimate the SD, as implemented in R’s `mad` function and with the default `constant` term). We also omitted individuals whose missingness rate is more than 6 SD from the median (likewise using R’s `mad` to estimate the SD).

To detect trios, we first located candidates as a prospective child that shares > 85% IBD1 and < 10% IBD2 to each of two prospective parents (IBD1 and IBD2 are the proportion of the genome in which IBD is shared on one haplotype and two haplotypes, respectively). We required that the prospective parents in any trio share ≤ 20% IBD1 with each other. Next we required that the two parents have different sexes and that the mother was ≥ 15 and ≤ 50 years old and the father was ≥ 15 and ≤ 80 years old when the child was born (using self-reported ages for each person).

From these inferred trios, we ascertained trio subjects for our analysis as those where each person is research-consented and typed on our v5 array, and we considered only one arbitrarily chosen child for any two parents. Next we determined monozygotic (MZ) twins or duplicates as sample pairs who share > 3,400 cM of IBD2; we then iteratively constructed sets in which each member is a MZ twin or duplicate of at least one other individual in the set. We arbitrarily chose one sample from each set to retain and excluded all the other samples from the trios, removing a trio if any member (parents or child) was among those to be excluded. This does not exclude trios with overlapping members, so a person can be a child in one trio and a parent in another or can have a trio child with multiple partners. Lastly, we used ancestry proportion estimates from 23andMe’s Ancestry Composition algorithm [11] to randomly ascertain 1,000 trios from the following six ancestry groups using the indicated criteria: Ashkenazi Jewish (> 95% Ashkenazi ancestry), European (> 95% European and < 30% Ashkenazi ancestry), admixed African (> 40% African, > 5% European, < 2% Indigenous American, and African + European ancestry > 95%), admixed American (> 20% Indigenous American, > 5% European, and American + European + African ancestry > 95%), East Asian (> 95% East Asian ancestry), and South Asian (> 95% South Asian ancestry). Note that the thresholds used do not in any way define these populations; this ascertainment is meant to provide a set of samples for evaluating performance across a range of ancestry backgrounds.

23andMe research participants provided informed consent and volunteered to participate in the research online, under a protocol approved by the external AAHRPP-accredited IRB, Ethical & Independent (E&I) Review Services. As of 2022, E&I Review Services is part of Salus IRB (https://www.versiticlinicaltrials.org/salusirb).

### UK Biobank dataset

We identified 6,270 parent-offspring pairs in the UK Biobank, defined as pairs who have an IBS0 < 0.001 (i.e., the proportion of the SNPs in which the pair are identical-by-state at zero of the two alleles is less than 0.001) and whose kinship coefficient is < 0.30. We oriented the parent-offspring pairs using year of birth and found 1,022 trios, confirming that parents were opposite sexes and that the parents were sufficiently unrelated (kinship coefficient < 0.0442). We performed quality control in PLINK v2.00a6LM [30] with the following filters: minor allele frequency `--maf` 0.0001, genotype missingness `–geno` 0.05, Hardy-Weinberg equilibrium `--hwe` 10^−9^. This left us with a total of 525,427 SNPs across the autosomes and the X-chromosome.

### 1000 Genomes Project-UK Biobank dataset

In the UK Biobank, 998 of the 1,022 trios are described as British or Irish. To understand intra-chromosomal phasing performance across different ancestries, we merged the UK Biobank genotype data with the 1000 Genomes Project (TGP) on chromosome 2. This provides an additional 594 trios, 487 of which belong to non-European ancestry groups. We used the TGP high-coverage whole-genome sequencing data [5], which comes filtered (Mendelian error rate ≤ 5%, genotype missingness < 5%, Hardy-Weinberg Equilibrium > 1 × 10^−10^ in at least one superpopulation, and minor allele count ≥ 2). The TGP data are aligned to build GRCh38 and the UK Biobank data provides coordinates from build GRCh37. We lifted the UK Biobank data over to build GRCh38 [22, 18] before merging the datasets and kept only SNPs present in both datasets (52,528 SNPs on chromosome 2). We ran PCA on the merged dataset to confirm the absence of batch effects (not shown). We refer to the merged dataset as UKB-TGP. We use admixed American to refer to the American (AMR) TGP samples for consistency with the 23andMe analysis. For non-European UKB samples, we attempted to match their self-reported ethnic background with the 23andMe ancestry labels; e.g., there are five UKB trios described as Indian, Pakistani, or Bangladeshi which we label as South Asian. This gave us trio data for five broad groups: African (which includes both European-admixed and non-admixed individuals), South Asian, East Asian, admixed American, and European (although note that for our analyses we show results for UK Biobank British trio children separately from the other European ancestry samples). For a breakdown of the sample composition of each ancestry group see Table S6.

### Assessing phase quality

We assess the performance of the phasing algorithms by comparing their output to the ‘true’ phase, which can be obtained with Mendelian phasing of the trio children and their parents. We used the Ancestry and Kinship Tools (AKT) software package [2] to perform Mendelian phasing; we refer to the output of AKT as the ‘trio-phased VCF’. The standard metric for phase quality is the switch error rate (SER), which is the proportion of consecutive heterozygous sites phased incorrectly (Figure 1A). Browning and Browning [3] found that most observed switch errors are consecutive switch errors, meaning a single SNP is phased incorrectly but the surrounding haplotype is phased correctly. This is penalized as two switch errors and inflates the SER. We refer to consecutive switch errors as ‘blips’ [26] and computed the SER with blips (where a blip counts as two switch errors, as is the standard) and without blips.

**Figure 1.**
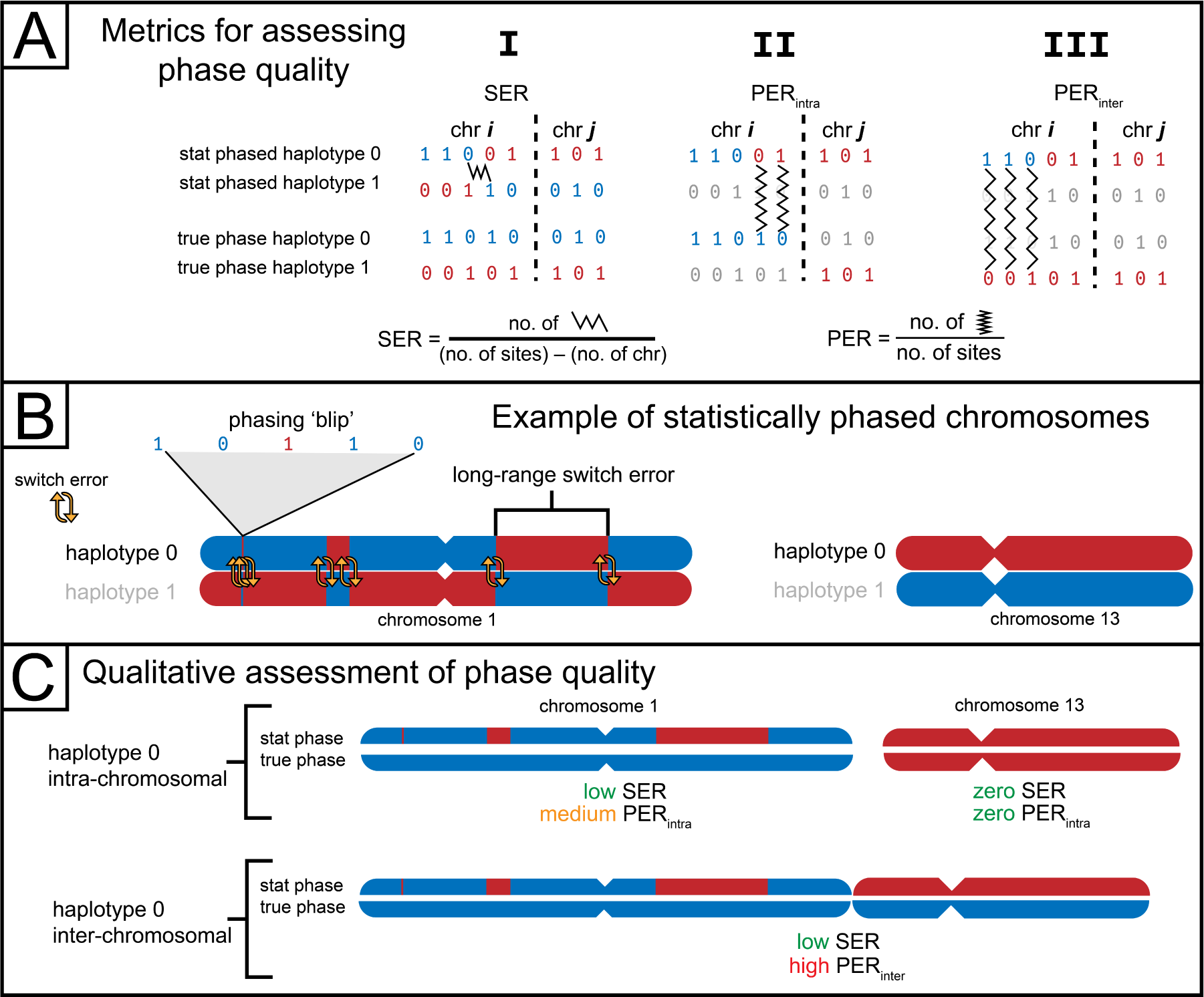
An overview of how we assess phase quality. **(A)** Three different metrics: switch error rate (SER), intra-chromosomal phase error rate (PER*_intra_*), and the inter-chromosomal phase error rate (PER*_inter_*). We show a sequence of eight heterozygous sites on two chromosomes. The top rows are the statistically-phased haplotypes and the bottom rows are the trio-phased (true) haplotypes; variants are colored by the known parent-of-origin (blue for paternal, red for maternal). The SER is depicted in **I** by a single switch error between the third and fourth positions, where the color of the statistically phased haplotype switches from blue to red. We show PER*_intra_* in **II** where we assess the proportion of mismatched sites between the statistical haplotype and the true haplotype. In this example, there are two regions since the markers fall on different chromosomes, so we would compute two different PER*_intra_*. Note that the two regions are compared to different true haplotypes that minimize PER*_intra_* (i.e., the first region is compared to the paternal haplotype and the second region compared to the maternal haplotype). In **III**, we show PER*_inter_*, which is computed for all regions against the same haplotype (in this case, the maternal haplotype). **(B)** Depicts two statistically-phased chromosomes. Chromosome 1 contains six switch errors, two of which involve a phasing blip. Another two involve a long-range switch error, where the parental haplotype is not corrected for a long genomic distance. We also show Chromosome 13 as being perfectly phased, noting that haplotype 0 largely corresponds to different parents between Chromosome 1 and Chromosome 13. **(C)** Shows, qualitatively, our three different phase quality metrics for the example in (B). There are two regions corresponding to Chromosome 1 and Chromosome 13. For chromosome 1, the SER is low (there are six total switch errors out of a possible ∼ 50k), but PER*_intra_* is moderately high due to the long-range switch error. For chromosome 13, both the SER and PER*_intra_* are zero because the chromosome is phased perfectly. However, PER*_inter_* is high because the two chromosomes’ first haplotype is compared to the same parental haplotype across both chromosomes.

We are also interested in assessing long-range phase quality in some genomic region. The region *r* can be an entire chromosome or a sub-region of a chromosome (the latter is important later; see **Inter-chromosomal phasing algorithm**). We call this metric the intra-chromosomal phase error rate PER*_intra_*, which we compute using the Hamming distances between the first inferred haplotype of the region and each of the two parental haplotypes. More formally, if *H_rP_*and *H_rM_* are the Hamming distances between the first haplotype of region *r* and the transmitted paternal and maternal haplotypes (respectively) and *n_r_* is the number of heterozygous sites in *r*, then

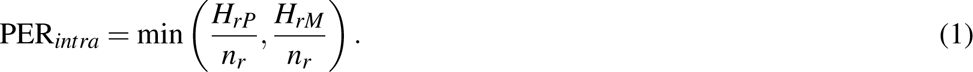

PER*_intra_* is a harsher metric than SER (see Figure 1C) because a single switch error will result in a small SER but can result in a large PER*_intra_*, depending on where the switch error occurs in the chromosome or region, e.g., a single switch error in the middle of a chromosome results in the highest possible PER*_intra_*, 0.50.

We are also interested in inter-chromosomal phase, in which the first haplotype always corresponds to the same parent both within and *between* chromosomes. To quantify inter-chromosomal phase quality, we define a metric similar to PER*_intra_*, called PER*_inter_*. The key difference is that, for all chromosomes, we compare the first inferred haplotype to the two trio-phased haplotypes, retaining the scores across chromosomes. (We ensure that the trio phaser orders the haplotypes such that the same parent, e.g., the father, transmitted the first trio-phased haplotype and the other parent transmitted the second haplotype.) More formally,

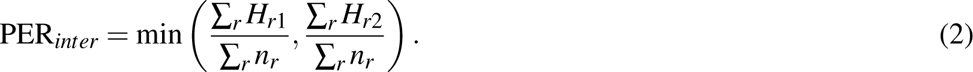

A more detailed explanation of these phase quality metrics is in Figure 1.

### Intra-chromosomal phasing

First we assessed the performance of two state-of-the-art phasing tools: SHAPEIT v5.1.1[19] and Beagle v5.4 [4]. We performed this analysis on chromosome 2 and computed the switch error rates (SERs) using trio-phased data as the true phase. We used the Ancestry and Kinship Tools (AKT) [2] to perform Mendelian phasing of the trios and the `--diff-switch-error` option of `vcftools` [9] to calculate SERs; this pipeline ignores sites with ambiguous trio phase (e.g., those heterozygous in all trio members), and AKT outputs unphased genotypes for these sites.

Most of our analyses were ‘out-of-sample’ runs (unless otherwise stated), where the trio children are phased against a previously-phased reference panel consisting of other samples (excluding the trio parents). This is in contrast to ‘in-sample’ runs, where the trio-children are jointly-phased with other samples (excluding the trio parents). A schematic of our phasing runs can be found in Figure S1. When phasing the 23andMe data, we divided chromosome 2 into overlapping regions of roughly 10 Mb each with an additional roughly 2 Mb overlapping the flanking regions (total window size ∼ 14 Mb). We then ligated the resulting phased regions using `bcftools concat -l`.

To collect runtimes, in the 23andMe data, we performed out-of-sample phasing by running both methods with 16 threads on two roughly 14 Mb regions and tracked wall-clock time and memory usage on either a C6I 32xlarge or R6I 32xlarge AWS ec2 instance. (We selected these regions as those that had median and 66.7-percentile switch error counts in a previous analysis with 8.1M samples and a different set of trios.) These AWS instances both have Intel Xeon 8375C (Ice Lake) processors with 128 vCPUs and either 256 GB (C6I) or 1,024 GB (R6I) of RAM. For Beagle, we previously converted the 8.6M data to `bref3` format and provided this as input; for SHAPEIT, we provided these data in BCF format. For the UKB-TGP, we used the computing resources of Brown University’s Center for Computation and Visualization. For the out-of-sample phasing runs, we ran both methods with 32 threads and 128GB of RAM using AuthenticAMD’s AMD EPYC 9684X 96-Core Processor (192 CPUs) and tracked wall-clock time. For the UKB-TGP, we provided data in the VCF format for all sites on chromosome 2 in a single run.

### Ancestry-specific phasing

We sought to investigate the phasing performance of non-European ancestry samples in the UKB-TGP joint dataset, which is 87% British-ancestry. We analyzed four broad, continental ancestry groups: African, South Asian, East Asian, admixed American, noting that most trios in these groups are from the TGP; see Table S6 for the non-British UKB samples included. For each ancestry group, we performed seven different in-sample runs (on chromosome 2) using both Beagle and SHAPEIT; each run contained the non-European ancestry samples (except for trio parents) and a random subset of European-ancestry samples (any EUR sample in TGP or British/Irish/European samples in UKB). Each run differed by the ratio of European samples to ancestry group samples: 1:8, 1:4, 1:2, 1:1, 2:1, 4:1, 8:1, similar to the analysis shown in Figure 4 of [33]. We also included a run containing only the ancestry group samples (i.e., without Europeans).

### Correlating IBD depth and switch errors

We also sought to understand the relationship of identity-by-descent (IBD) sharing and switch errors, hypothesizing that, while SHAPEIT and Beagle do not use IBD explicitly, they do phase using individuals with similar haplotypes to a focal sample. Therefore, individuals who share more IBD with other samples are likely to have lower SERs, particularly in regions covered by IBD. To quantify per-locus IBD rates, we sought idealized phase for the trio children. To that end, we used the trio-phased VCF as a “scaffold”, passing this and a reference panel containing the non-trio UKB-TGP individuals (who were phased in-sample) as input into SHAPEIT [19]. This scaffold-based inference phases any unphased sites in the trio VCF (i.e., those that could not be phased due to Mendelian logic, including those where all trio members are heterozygous). We used the resulting data to call IBD against the remaining UKB-TGP dataset using `TPBWT` [13]. For each focal individual, we computed the number of IBD segments that cover each site on chromosome 2. We also know which sites flank a switch error (from the `vcftools` output) and merging these two datasets gives us, for each site and each focal individual, the number of IBD segments shared with the focal individual that cover it and a boolean indicating whether the site belongs to a switch error.

### Inter-chromosomal phasing

#### Pre-phasing step

Our inter-chromosomal phasing method, HAPTIC, takes as input a file containing IBD segments generated from the haplotypes output by a ‘pre-phasing’ step. This pre-phasing step differs for the UKB and 23andMe due to the computational constraints of phasing in the 8.6M 23andMe dataset. We first describe this step in UKB, where we performed in-sample phasing with Beagle run on the entire dataset, including the trio children but excluding their parents. This ensures that the phase quality of trio children is not biased by the presence of their parents. We performed additional in-sample phasing of the entire dataset, this time including the trio parents and excluding the trio children; the trio parents were subsetted from this output and merged with the output of the first pre-phasing run. While this second step increases the comprehensiveness of IBD detection, its impact on the overall results is expected to be minimal, affecting only about 0.60% of pairs. Due to computational limitations, only the first pre-phasing step (excluding parents) was performed for the 23andMe dataset.

Next we outline the pre-phasing step in the 23andMe dataset. As noted earlier, we performed in-sample phasing of the 8.6M 23andMe research subjects on chromosome 2 while excluding the trio parents. However, our SHAPEIT phasing runs for the other chromosomes included the parents and consequently have higher phase quality for the trio children than observed in the chromosome 2 data (not shown). We used an approach to mimic phasing the children with the parents excluded genome-wide as follows. First, we detected switch errors in the original phase (i.e., that with the parents included), corrected these errors (but retained all original blips), and then randomly added new non-blip switch errors at the same rate observed for each child in the chromosome 2 data (if the child had 0 switch errors, we set the rate to 10^−6^). Introducing new switch errors proceeded by randomly sampling a Poisson number of switches to add given the chromosome 2 rate and the number of heterozygous sites on the chromosome being corrupted; we then sampled switches by weighting each heterozygous site by its cM genetic distance from the previous heterozygous site and assigning 0 probability to the first and last heterozygous sites (as switch errors necessarily occur between two heterozygous sites). This approach therefore uses a genetic map-based probability of sampling a switch error. We used the HapMap genetic map on GRCh38 coordinates available from the Beagle 5.4 website for cM positions in this step. This map encompasses nearly all SNPs we analyzed, but we set the genetic position to 0 or the last cM position on a chromosome for sites with physical positions respectively before or after those included in the map. This affected 15 SNPs or fewer on the autosomes, but the X chromosome includes > 500 positions outside this genetic map. To get cM positions for these sites, we extrapolated the chromosome X map at a rate of 1.18 cM/Mb (its chromosome-wide recombination rate) to sites not included in this map.

When `vcftools` detects switch errors, it outputs a start and end position of each error—the heterozygous sites that delineate the phase switch. Often the intervening SNPs are homozygous in the target individual, but AKT retains unknown phase at Mendelian-ambiguous positions and a switch error can span such sites. This is relevant to correcting the original switch errors: we randomly chose a position between the switch error’s start and end positions to begin inverting the phase, and this could introduce a small number of undetected switch errors at the Mendelian-ambiguous heterozygous sites.

After the pre-phasing step, we computed identity-by-descent (IBD) segments ≥ 5cM using `TPBWT` [13]. All parameters were default values, except we set `use_phase_correction` to `False`. Here HAPTIC differs markedly from [29]: we start with IBD computed from phased data and seek to correct long-range phase errors and infer inter-chromosomal phase.

#### Inter-chromosomal phasing algorithm

We now describe HAPTIC from the perspective of a single focal individual *F*. Our goal is to group the IBD segments shared with *F* into two clusters corresponding to maternally- and paternally-inherited IBD segments, although in practice the parent-of-origin (i.e., the parental sex) of the clusters is often unknown (it can sometimes be inferred for males using X-chromosome IBD sharing as done in [20]). We start by taking IBD segments shared between *F* and their relatives and create IBD ‘tiles’: a set of IBD segments that overlap by ≥ 3cM on either haplotype. (So, for example, two non-overlapping 5 cM segments on haplotype 1 can be joined in a tile using a segment that overlaps both of them on the other haplotype by 3 cM.) We define a sub-tile as the IBD segments in a tile that fall on the same (pre-phased) haplotype.

Next, we construct a signed graph *G*; nodes in *G* correspond to IBD segments. We use the procedure below to generate edges, seeking to add positively-weighted edges between two IBD segments inherited from the same parent of *F*, and, conversely, negatively-weighted edges between two IBD segments inherited from different parents of *F*. A node *n_i_* has several attributes that are relevant: *r* is the ID of the relative who shares the IBD segment with *F*; *l* is the length of the corresponding IBD segment; *k* is the sum of all IBD (in cM) that *F* and *r* share; *t* is the tile the IBD segment falls in; *s* is the sub-tile the IBD segment falls in (*s ∈ {*0, 1}; this is also the haplotype index the IBD segment falls on); lastly, *c ∈ {*0, 1} is the parent index, which is unknown. We denote the attribute value of a node as *n_i_*(*a*) where *a* is the attribute; e.g., *n_i_*(*l*) is the length of the IBD segment corresponding to the *i*th node. In the special case in which two IBD segments shared between *F* and the same relative are adjacent on the haplotype and separated by a gap of ≤ 1cM, we merge the two IBD segments and assign them to the same node, where *n_i_*(*l*) is the sum of the two segment lengths. Importantly, if the two IBD segments belong to different tiles, this does not merge the tiles (although the node can be found in two tiles).

We add edges to *G* in two conditions. First, we add edges between nodes *n_i_* and *n _j_* when *n_i_*(*t*) = *n _j_*(*t*) (i.e., the nodes belong to the same tile). We assume (and confirm in Figure 7) that the phase quality is high in tiles, with a small number of phase errors resulting from genotype errors and/or blips and the coarseness of IBD segment ends. Under this assumption, IBD segments on the same haplotype should be inherited from the same parent and IBD segments on opposite haplotypes inherited from different parents (assuming the IBD segments are in the same tile). We represent this “sidedness” with the sign of the edge weights: positively-weighted edges connect nodes (IBD segments) inherited through the same parent and negatively-weighted edges connect nodes inherited through different parents. The edge weight *e_i j_* for nodes in the same tile is given by:

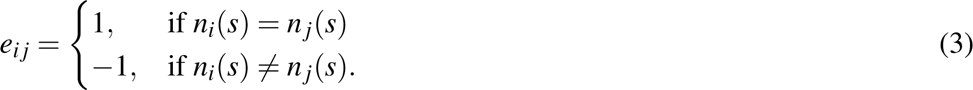

Second, we also add edges between nodes if *n_i_*(*r*) = *n _j_*(*r*) (i.e., their corresponding IBD segments are shared with the same relative) and if certain length criteria are satisfied. To do this, we define two thresholds: *t*_1_ and *t*_2_. The edge weight *e_i j_* is given by:

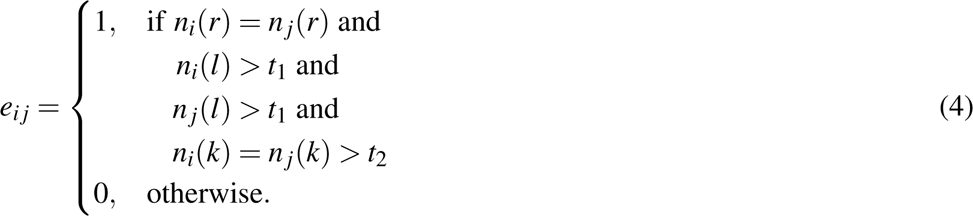

This leverages the assumption that a pair of IBD segments shared with the same relative is uniparentally inherited. However, we find frequent violations of this, even among non-descendant relatives, particularly when the IBD segments are short in length or if *r* is distantly related to *F*. This motivates the *t*_1_ and *t*_2_ thresholds: edges are only added between pairs of IBD segments shared with the same relative if the segments are both sufficiently long (*n*(*l*) > *t*_1_) and the relative has sufficient total IBD sharing (*n*(*k*) > *t*_2_). The result is a signed graph *G* = (*V, E, A*) where *V* are vertices (these are the IBD segments, which we refer to as nodes), *E* are the edges, and *A* is the adjacency matrix. Having constructed a signed graph, our goal is to find two partitions of *V* : *X* and *Y* = *V\X*, where nodes in *X* are IBD segments inherited from one parent and nodes in *Y* are IBD segments inherited from the other. To that end, we draw from the work of [24] who extended bipartite spectral clustering to signed graphs. They use the signed Laplacian matrix as 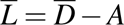, where 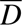 is the signed degree matrix, a diagonal matrix given by:

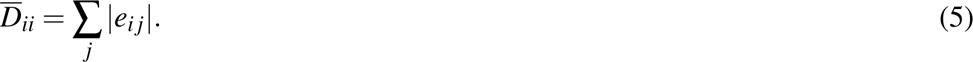

We compute the eigenvectors and eigenvalues of 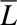 using NumPy’s `linalg.eigh` function [17]. The eigenvalues (and their corresponding eigenvectors **v**^(1)^, **v**^(2)^*, …,* **v**^(^*^n^*^)^) are sorted, such that *λ*_1_ < *λ*_2_ < … < *λ_n_*, where *n* is the number of nodes in *G*. Bipartite clustering is performed using **v**^(1)^. We infer the parent index of a node *n_i_*(*c*) as:

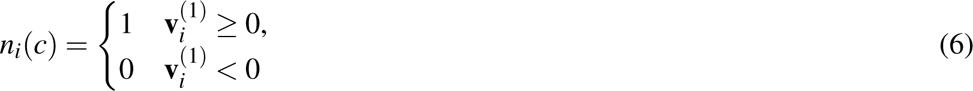

Briefly, this procedure is an approximation of the minimum signed ratio-cut problem [24]. The signed cut of the graph, denoted scut(*X,Y*) is the number of edges that are (1) positively-weighted and connect nodes between cluster *X* and *Y* or (2) negatively-weighted and connect nodes within *X* or within *Y*. The signed ratio cut weights scut(*X,Y*) by 1*/|X|* + 1*/|Y|*, thus favoring cuts of *V* that are somewhat balanced in size and preventing cuts in which *X* or *Y* have a small number of vertices [24]. Minimizing the signed-ratio cut is NP-hard because the desired output are discretized cluster labels [32]. By relaxing the requirement that cluster labels are discretized, the eigenvector associated with the smallest eigenvalue can be used to approximate the minimum signed-ratio cut [24]. However, we must map the eigenvector to discretized cluster labels, which we do using Equation 6 [24].

In practice, we use the largest connected component (in terms of the number of nodes) of *G*, *G*_1_, to construct 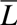. Different components of *G* are formed of disjoint sets of tiles, meaning that the underlying IBD segments in one component are shared with a set of relatives that do not overlap those of another component. (Note this is not strictly true since IBD segments *i* and *j* shared with *r* could be in two components if *n_i_*(*l*) < *t*_1_ or *n _j_*(*l*) < *t*_1_ or *n_i_*(*k*) < *t*_2_.) We cannot resolve the phase between two components of the graph, so we use only the largest component of the graph for phasing. This means, unless there is only one component that overlaps all sites, HAPTIC will not phase all the variants.

If the graph is balanced, then *λ*_1_ = 0 [24]. A balanced graph indicates that there are no positively-weighted edges between clusters, nor are there negatively-weighted edges within a cluster. With respect to our task, a balanced graph also indicates that there are no logical inconsistencies in our graph. For example, if three nodes are all connected by negatively-weighted edges, this is logically inconsistent (and impossible) for IBD transmitted from two parents. Importantly, *λ*_1_ = 0 does not necessarily indicate that the clustering is perfect, and graphs where *λ*_1_ > 0 are not necessarily clustered imperfectly. However, we found empirically that graphs where *λ*_1_ *≫* 0 are likely poorly clustered (not shown).

After the IBD segments are clustered, the final step is to correct switch errors between tiles and perform the inter-chromosomal phasing. Recall that *n_i_*(*c*) is the inferred parental label of node *n_i_*, which belongs in tile *n_i_*(*t*) and sub-tile *n_i_*(*s*). The set of (tile, sub-tile) tuples for parent 1 is {(*n_i_*(*t*)*, n_i_*(*s*))|*n_i_*(*c*) = 0} and for parent 2 is {(*n_i_*(*t*)*, n_i_*(*s*))*|n_i_*(*c*) = 1}. Ideally, the intersection of these sets is the empty set. A non-empty set indicates that IBD segments from the same sub-tile were clustered in different parental clusters; in these cases, the entire tile is removed. Further, if the (tile, sub-tile) tuples (*t,* 0) and (*t,* 1) are in the same set, the entire tile is removed (in this case, IBD segments from different sub-tiles in the same tile were clustered in the same parental cluster). To phase the variants, we arbitrarily decide that the parent corresponding to the first (tile, sub-tile) set will be the first haplotype. If this is (*t,* 0), the haplotype phase is not switched (recall that *n_i_*(*s*) corresponds to the haplotype index of the input phase). If this is (*t,* 1), the haplotype phase is switched.

#### Identifying descendants

Including IBD segments from a descendant *d* of focal individual *F* leads to poor clustering because the IBD segments they share will be on both parental haplotypes of *F*; thus, identifying and removing descendants is a key component of HAPTIC. Descendants of any full-siblings of *F* cause the same problem, this when we refer to “descendants” we are referring to any descendants of *both* of *F*’s parents. Our approach uses a threshold for the degree of relatedness of potential descendants. For instance, if we do not expect the dataset to span more than three generations, the most distant descendant of *F* would be *F*’s full-sibling’s grandchild, who is a third degree relative of *F*. If we set our descendant threshold as third degree, then all relatives who are second or third degree are referred to as potential descendants, which we denote 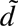. Note that first degree relatives, including full-siblings and offspring of *F* are always excluded by our inter-chromosomal phasing algorithm.

We seek to determine which of the potential descendants are true descendants and which are not: such close non-descendant relatives are a rich source of information for inter-chromosomal-phasing. To that end, we apply the inter-chromosomal phasing algorithm, but exclude potential descendants (i.e., second and third degree relatives) from the analysis. The result is a set of genomic coordinates and haplotype indices that correspond to each of *F*’s parents. For most focal individuals, these genomic coordinates will not cover the entire genome, so there will be regions where the parental phase is unknown. We expect that a descendant of *F* will share 50% of IBD with *F* on *F*’s maternal haplotype and 50% of IBD with *F* on *F*’s paternal haplotype. Of course, there will be variance around the 50% expectation due to meiosis (and this variance will increase with the distance of the relative) and because the parental phase may only cover a small portion of the genome. Next, for each 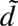, we take all the IBD segments 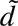 and *F* share and compute the total overlap (if any) of the IBD segments with each parent’s haplotypes. We label 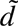 as a descendant if they share IBD with both parents of *F* and a non-descendant otherwise. We consider 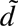 to share IBD with both parents of *F* if at least 20% of the IBD that we are able to assign in this step is shared with both parents. Descendants’ IBD is excluded from the input IBD and the algorithm is rerun.

Our method of finding descendants has several advantages over other methods: [29] and [20] identify descendants as individuals who are closely related to a pair of close relatives of *F*, *r*_1_ and *r*_2_, where *r*_1_ and *r*_2_ are unrelated. (For example, if *F* has a maternal aunt, a paternal aunt, and a nephew, we can identify the nephew as a descendant because he will be related to both aunts of *F* but the aunts will typically not be related to each other.) However, this approach will not work if (1) there is only one other close relative; or (2) there are only close relatives on one side of the family. If close relatives are equally likely to belong to either side of the family, this would require ≥ 6 close relatives in order to have a ≥ 95% chance of two of those relatives being on opposite sides of the family; moreover, relatives on each side of a family may be genotyped at unequal rates.

Because the UK Biobank has a relatively narrow age range, we do not expect many descendants, and any descendants that we see would likely be limited to nieces/nephews of *F*. The trio children in particular are extremely unlikely to have descendants, and indeed we found no relatives of a trio child who are fourth degree related or closer to both parents. To test our approach to identify descendants, we simulated both nieces/nephews and grandchildren of the trio children using Ped-sim [6]. To simulate a niece of *F*, we took the Beagle-phased VCF containing *F*’s parents and simulated grandchildren of those parents, allowing Ped-sim to randomly select a partner for the simulated aunt/uncle in the second generation. We then called IBD between the Beagle-phased VCF of *F* and the newly simulated niece/nephew. (Note we retained the ‘true’ phase of all nieces/nephews/grandchildren as outputted by Ped-sim). To simulate the grandchild of *F*, we input *F*’s trio-phased VCF into Ped-sim and simulated *F*’s grandchild, again allowing Ped-sim to randomly sample partners. IBD between *F* and their grandchild was then called using the Beagle-phased VCF containing *F*, which allows us to merge the IBD segments with the IBD segments shared between *F* and their real relatives. We performed this simulation for each UK Biobank trio child, ran our inter-chromosomal phasing descendant detector, and kept track of true positives (simulated descendants that we identified as such) and false positives (non-descendants identified as descendants) as well as the total tile coverage (in cM) of the phasing algorithm run without potential descendants.

## RESULTS

### Intra-chromosomal phasing

Our intra-chromosomal out-of-sample phasing of 23andMe data show remarkably low SER, with the lowest SER achieved for admixed African individuals phased with Beagle: 0.012% (sd 0.035%) (Figure 2, left). Here, we focus on the blip-removed phasing runs as we are generally more concerned with long-range haplotype phasing and because past work has shown that blips may be primarily due to genotyping error [3]. Unless otherwise noted, all figures are Beagle results (SHAPEIT results can be found in the Supplement). The highest mean SER in 23andMe was found in South Asian ancestry individuals phased with SHAPEIT (mean 0.68%, sd 0.62%) (Figure S3, left).

**Figure 2.**
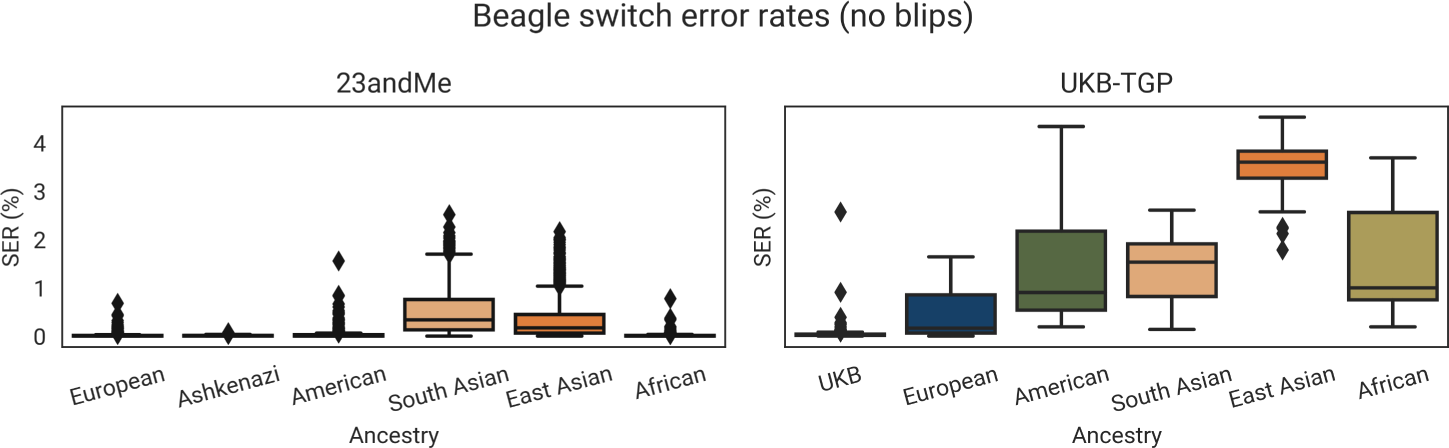
Beagle switch error rates (SER) excluding phasing ‘blips’ for 23andMe trio children (left) and UKB-TGP trio children (right), split by ancestry group.

A general trend we see is that 23andMe European, Ashkenazi, admixed African, and admixed American ancestry individuals tend to have low and comparable SER, i.e., the range in SER from Beagle is from 0.012% (admixed African, sd 0.03%) to 0.028% (admixed American, sd 0.08%) (Figure 2, left). Mean SER in South and East Asians are higher: 0.36% (sd 0.17%) for East Asians and 0.50% (sd 0.34%) for South Asians.

We also note that the variance in SER is much higher for the Asian ancestry groups. A starker difference between the non-Asian and Asian groups can be seen in the distribution of the number of switch errors. Most non-Asian individuals have zero switch errors on chromosome 2 (ranging from 45.1% of Ashkenazi individuals to 57.4% of admixed African ancestry individuals), whereas only 2.9% and 7.0% of South and East Asians, respectively, have zero switch errors (Figure 4).

Beagle generally has lower SER than SHAPEIT (Figure 3). However, this performance varies between the non-Asian and Asian ancestry groups. Of the non-Asian ancestry groups, the median per-sample difference between SHAPEIT SER and Beagle SER is zero and the mean difference is small, e.g., 0.0012% for admixed African ancestry individuals and 0.00086% for admixed American individuals (Table S4). However, for the Asian ancestry groups, the mean differences are 0.18% for East Asians (sd 0.226%, median 0.10%) and 0.177% for South Asians (sd 0.21%, median 0.21%). Interestingly, the difference in SER is higher when computed *with* blips (Table S3), suggesting that SHAPEIT produces more blips than Beagle. Despite the SER being, on average, lower for Beagle-phased samples, the number of samples with zero switch errors is actually higher for SHAPEIT-phased European, admixed American, South Asian, and admixed African ancestry individuals, e.g., 57.4% of Beagle-phased admixed African ancestry individuals have zero switch errors (Figure 4) compared to 69.3% SHAPEIT-phased individuals (Figure S7). Even with the general trend of SHAPEIT phasing *more* individuals with zero switch errors, SHAPEIT also phases more individuals with 6+ switch errors than Beagle.

**Figure 3.**
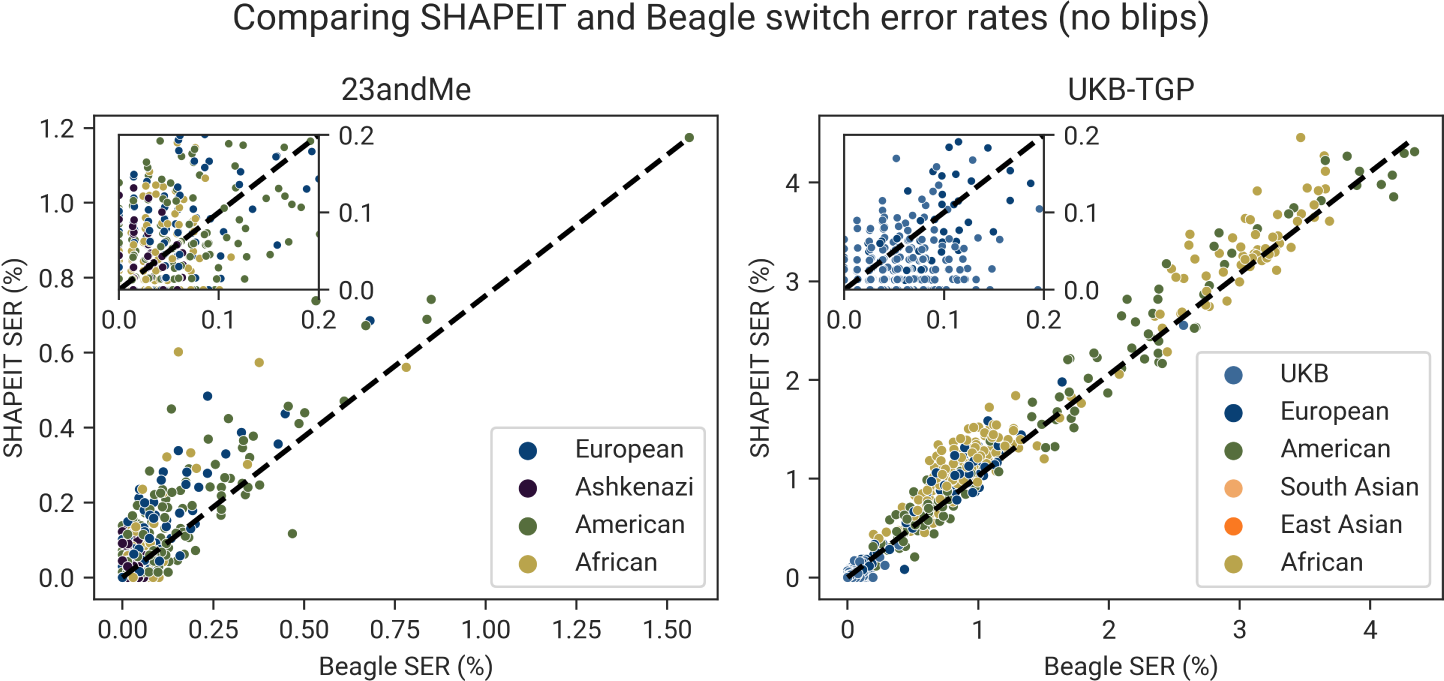
Comparison of Beagle switch error rates (SER; on x-axis) and SHAPEIT SER (y-axis) with phasing blips removed. Each point is a trio child, colored by their ancestry group (note that this does not include South and East Asian trio children (see Figure S6 right panel). The dashed line indicates an equal Beagle SER and SHAPEIT SER. As most 23andMe trio children have SER < 0.20%, we included an inset showing these trio children.

**Figure 4.**
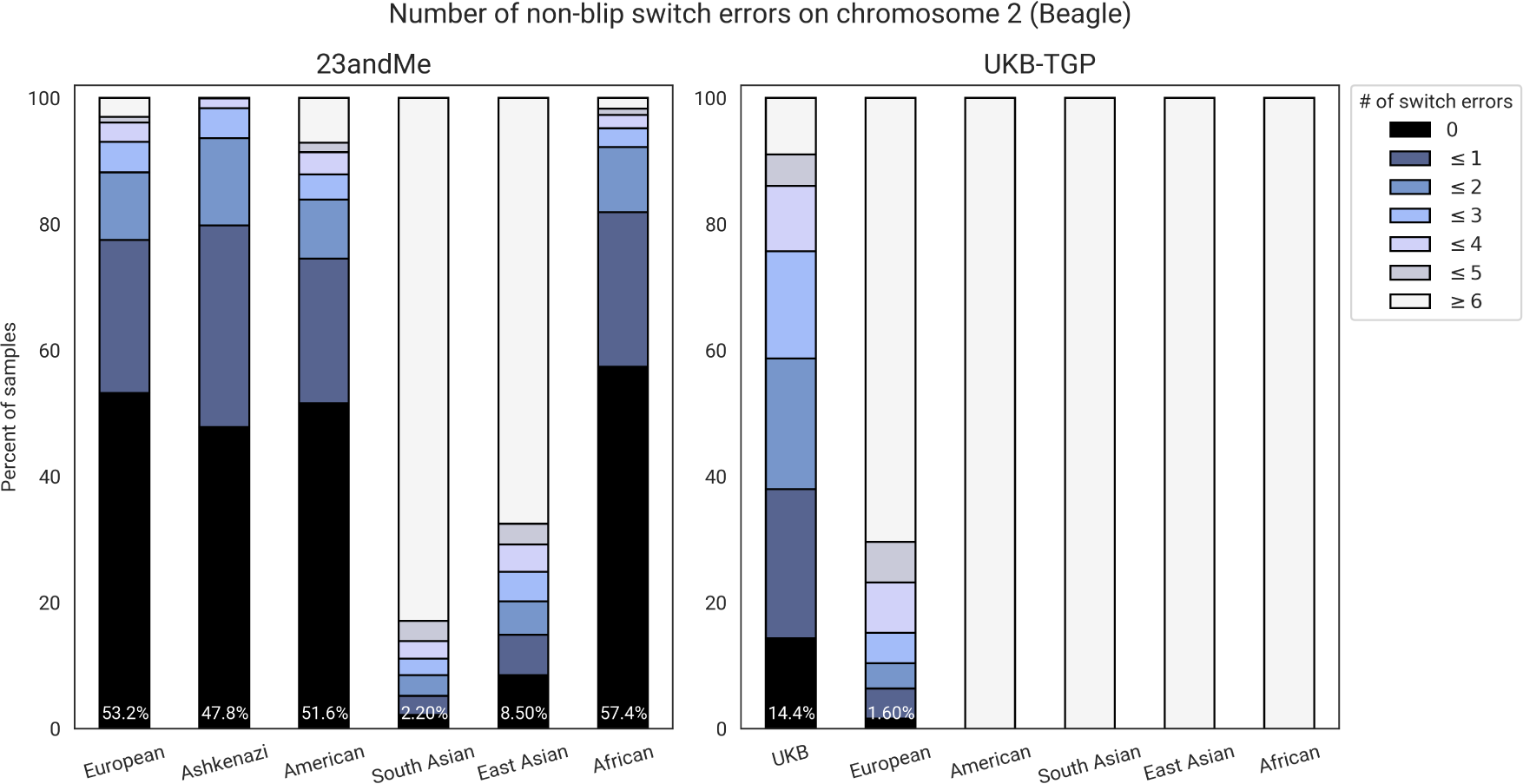
A stacked bar plot showing the distribution of the number of switch errors (Beagle phasing; blips removed) in the trio children, split by dataset and ancestry group. For example, 53.2% of 23andMe European-ancestry trio children have zero non-blip switch errors. On the other hand, all admixed American, South Asian, East Asian, and African UKB-TGP trio children have ≥6 non-blip switch errors.

Next we looked at the performance of Beagle and SHAPEIT on the UKB-TGP dataset (Figure 2, right). Here, most non-European ancestry trio children are from 1000 Genomes Project (TGP). The combined dataset of UKB-TGP is still overwhelmingly white British or Irish (90.3%; an additional 3.3% are continental European), and so we do not expect comparable SER to 23andMe for the non-European ancestry trio children. Indeed, we find the biggest difference between the two datasets in East Asian individuals: UKB-TGP East Asian trio children have, on average, 9.6x more switch errors than East Asian trio children in 23andMe, with a mean SER of 3.49% (sd 0.54%) (Table S2). The lowest SER in this merged dataset is found in the white British UKB trio children with a mean SER of 0.021% (sd 0.09%); note that that is remarkably close to the mean 23andMe SER for European individuals of 0.0172%. For the non-British European TGP samples (CEU and IBS), the mean SER was 0.43%, which is on par with the worst performing 23andMe groups (East and South Asians). All non-European groups have mean SER > 1%.

We kept track of the wall-clock time for SHAPEIT and Beagle in both the 23andMe and UKB-TGP phasing runs. Overall, we found SHAPEIT to be significantly faster. In the UK Biobank, the run time was 10.8 hours for Beagle versus 6.9 hours for SHAPEIT (∼36% faster) for chromosome 2. The 23andMe 8.6M dataset chromosome 2 run was split into ∼14 Mb regions (Methods). The average region runtime was 3,186 seconds (SHAPEIT) and 6,441 seconds (Beagle). With 278 autosomal regions, the estimated genome-wide runtime is ∼ 497 hours for Beagle and ∼ 246 hours for SHAPEIT (about 51% faster).

When comparing SHAPEIT and Beagle SER, we found that SHAPEIT performed slightly better than Beagle for white British UKB trio children (mean SER difference, SHAPEIT minus Beagle, −0.015%, sd 0.028%, median −0.013%), which is in contrast to all 23andMe ancestry groups and the remaining UKB-TGP groups (Table S4). For admixed American, African, and European ancestry UKB-TGP, Beagle outperforms SHAPEIT by a sizeable margin relative to its outperformance in the 23andMe groups, e.g., for Africans the mean difference is 0.23% in the UKB-TGP but 0.0012% in 23andMe. For East and South Asian groups, the mean difference in SER is comparable between 23andMe and UKB-TGP. For example, the mean differences for South Asians are 0.17% (23andMe) and 0.20% (TGP-UKB), and the mean difference for European trios is 0.006% (23andMe) and 0.065% (UKB-TGP).

The 1000 Genomes Project data allows us to further breakdown the results by more fine-grained population labels, e.g., some African (AFR) samples are continental African (GWD, ESN, YRI, LWK, MSL) and some are admixed with substantial European ancestry (ASW, ACB). As an example, the overall SER is higher for the African samples than the South Asian samples, which is in contrast to the 23andMe samples where the SER is much lower in the admixed African ancestry individuals compared to the South Asian ancestry individuals. However, when we split the samples by continental African and admixed African individuals, Figure S12 shows that SER is now twice as low for the admixed African samples (ASW, ACB) as the South Asian samples. For the continental African groups, we see there is a difference in SER that correlates with their representation in the UK Biobank. For example, ESN and YRI are Nigerian ethnic groups and there are 1,159 Nigerian-born individuals in the UKB (which is the fourth highest African born group in the dataset). On the other hand, GWD (Gambian) have the highest SER and are less represented in the UKB (with only 43 Gambian-born UKB participants). As another example, the SER is higher in Mexican (MXL) and Peruvian (PEL) samples than in Colombian (CLM) and Puerto Rican (PUR) samples; the latter two have significantly higher proportions of European ancestry than the former two [34]. We also note that the UKB-TGP admixed American ancestry group is by far the smallest group, with a total of 291 individuals, but has a comparable mean SER to South Asians (total of 8643 individuals); however, this is unsurprising given admixture, e.g., the median European ancestry proportion of CLM (Colombians) is 65% [34] and so these samples benefit from the European ancestry samples in UKB-TGP.

#### Ancestry-specific phasing

For all non-European UKB-TGP ancestry groups, we performed several in-sample phasing runs each with a different ratio of European-Group samples (where Group is an ancestry group). We compared the SER of eight different ratios of European:Group to the baseline SER computed from the in-sample phasing run with the entire UKB-TGP dataset (Figure 5). For admixed American trio children, the SER decreases as more European ancestry samples are added, and the lowest SER is found when they are run with the entire UKB-TGP merged dataset. Admixed American trio children see large changes in mean SER when more European samples are added: 2.32% when no European samples are included to 1.51% with the entire dataset included. This pattern is consistent between Beagle and SHAPEIT (Figure S8). Less consistent between SHAPEIT and Beagle are the patterns in South Asians, most which benefit from a small number of European samples relative to the South Asian sample size, but this differs depending on the population and the phasing algorithm. BEB (Bengali’s), for instance, find their lowest SER when no European samples are added (SHAPEIT) or when they are added at a ratio of 1:4 (Beagle). Otherwise, the best performing ratio ranges from 1:8 to 1:1. Beagle phases the East Asian trio children best at a European:East Asian ratio of 1:4; SHAPEIT phases the Han Chinese (CHS) best when no European samples are added and Vietnamese (KHV) at a ratio of 1:4.

**Figure 5.**
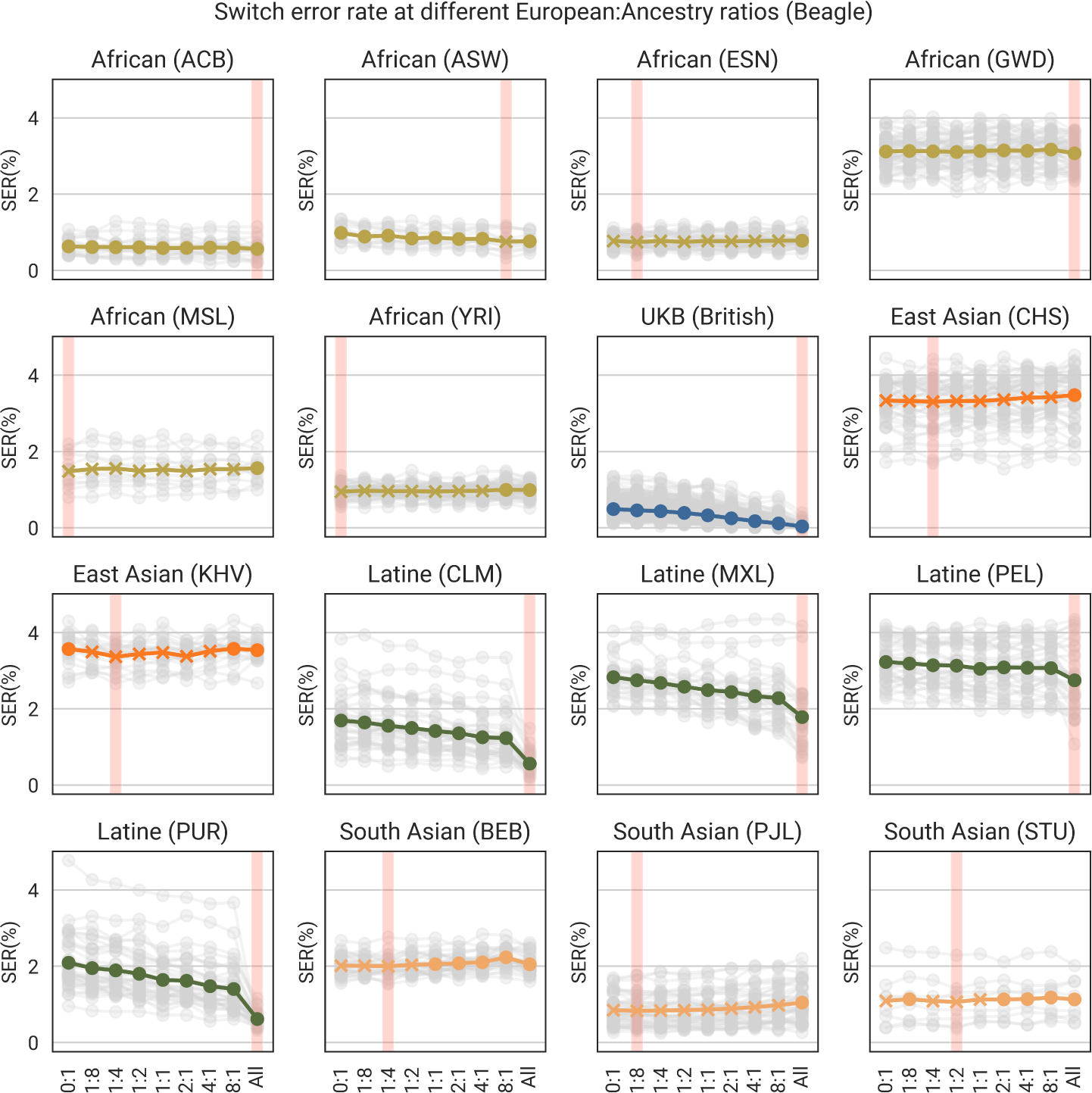
For each continental ancestry group, we did several Beagle ‘in-sample’ phasing runs where we varied the number of Europeans in the dataset relative to the size of the ancestry group. For instance, the admixed African 1:8 phasing run indicates that there was 1 European sample for every 8 African-ancestry samples. Note that for the UKB (British) samples, we included the trios with a random subset of British individuals to make a subset of 8000 the (“British-only” subset); a 1:8 ratio dataset would include an additional 1000 British/European samples for a total of 9000 individuals. These ratios are on the x-axis (‘All’ is the in-sample run with the entire dataset). Here, we show the results split by 1000 Genomes Project population labels. Each trio child’s SER are shown in grey, and the colored line is the mean line. ‘X’ markers indicate that the mean is below the ‘All’ (in-sample, entire dataset) mean. The red vertical bar indicates the ratio with the lowest mean SER for the population.

SHAPEIT and Beagle are fairly consistent when phasing the different admixed African ancestry groups. For instance, both phase the admixed African Americans (ASW) best at a European:admixed African ratio of 1:8 and most continental admixed African populations at ratios between 1:8 and 1:2. A notable exception is the difference in SHAPEIT and Beagle in phasing the Gambian (GWD) trio children. SHAPEIT’s lowest SER is achieved when there are no European ancestry samples added, whereas Beagle’s lowest SER is found when all samples are included. We note, however, that generally the range in SER for admixed African ancestry samples is small and we only ran one phasing run. As the phasing process is stochastic, it is unclear if the differences in SER here are statistically significant.

#### Identity-by-descent decreases switch error rates

We also sought to understand the relationship between identity-by-descent (IBD) sharing and switch error rates. We tested the hypothesis that SER should be generally lower in individuals with more IBD in the dataset and that switch errors occur in regions with lower IBD coverage. We computed IBD using the trio-phased VCF for the trio children and called IBD against the in-sample Beagle phasing run, which included all individuals in the UK Biobank and TGP (minus trio parents; Methods). This ensures that the IBD segments called are not affected by switch errors (at least, from the perspective of the trio children; switch errors may still occur in their relatives, which may artificially shrink or break up IBD segments). For each focal individual, we focus on the number of sites on chromosome 2 that are covered by IBD segments > 3cM. Figure S9 shows the distribution of the number of sites covered by IBD by TGP population, as well as the distribution for white British UKB trio children. The IBD coverage varies less between the broad continental groups and more between the finer-grained population groups. We also show the relationship between IBD coverage and SER in Figure S10, where we see that SER generally decreases with increased IBD sharing. Figure 6 shows the mean IBD coverage and mean SER for each population group in TGP and each self-reported ethnicity group in UKB. Again, we observe that groups with higher mean IBD coverage also have lower mean SER. The plot also illustrates the issues of continental or broadly-defined population-labels, as we see major differences in performance between finer-grained population groups within the same continental group, e.g., PJL (Punjabi’s) versus BEB (Bengali’s).

**Figure 6.**
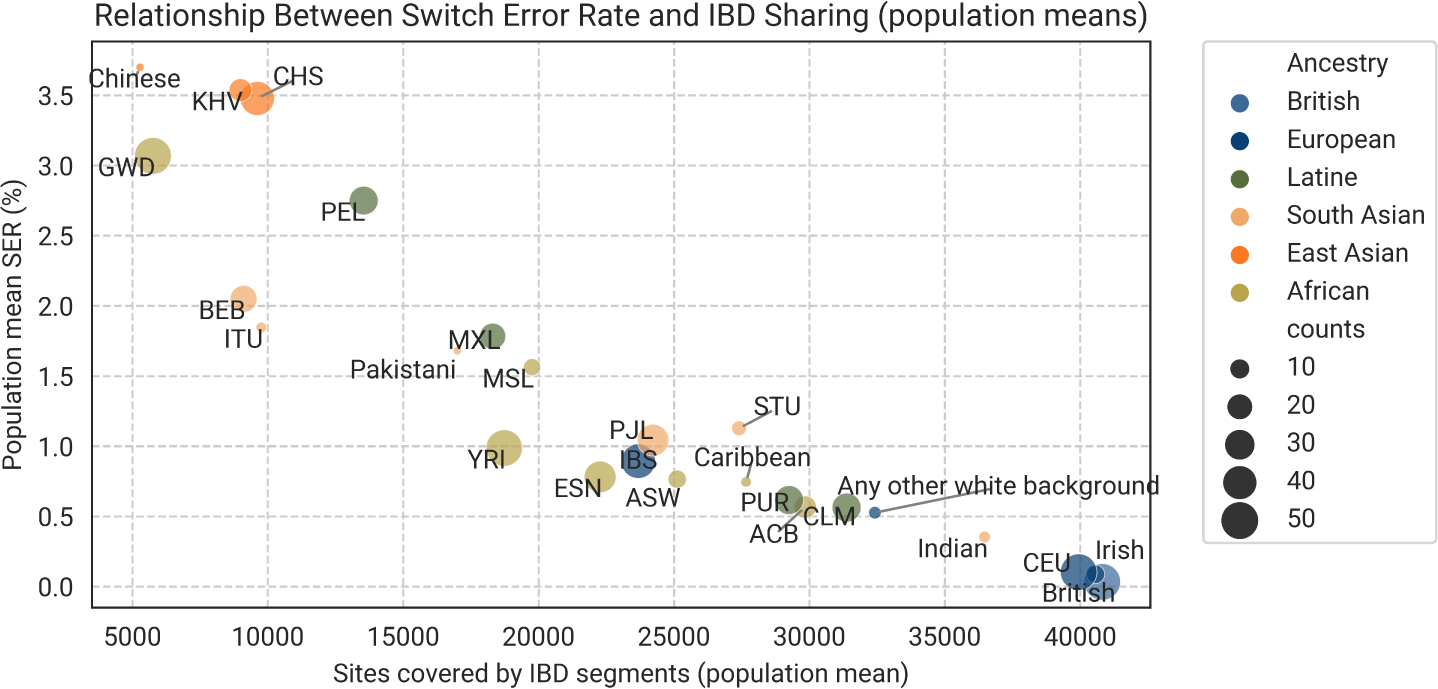
The relationship between switch error rate (SER) and IBD sharing. For each trio child, we computed the number of sites on chromosome 2 covered by at least one IBD segment and compare to the SER. We computed the IBD using trio-phased (Mendelian phasing with SHAPEIT scaffold phasing to phase unresolved triple-heterozygous sites [19]) genotype data. Here we plot the mean for IBD coverage and SER for each population (three-letter codes are TGP and other labels are UKB self-reported ethnicity labels). The size of the markers is proportional to the number of trios included, except for ‘British’ which we scaled to be the same size as the second largest group, CEU (there are 985 British trios).

Next, we analyze IBD sharing across switch errors. For each non-blip switch error, we calculated the mean IBD coverage over the switch error window (the window may be as small as two adjacent variants but can be much larger if the two heterozygous sites are separated by many homozygous or heterozygous sites that could not be resolved with Mendelian phasing). We then sampled, for each switch error window, 100 windows (of the same number of SNPs) that did not contain a switch error and computed the mean IBD coverage of each window. For each switch error, we now have the IBD coverage of the switch error and a genome-wide average of IBD coverage (IBD segments shared with the trio child) of the same-sized window. Figure S11 shows the results of this analysis split by ancestry group. For all ancestry groups, switch errors tend to be covered by less IBD (Mann-Whitney test *p <* 10^−8^ for all groups). For example, in the white British UKB samples, non-blip switch errors are exceedingly rare, but of the 440 non-blip switch errors, 330 occurred in windows covered by zero IBD segments, whereas the average window of the same size was covered by 23.9 IBD segments (sd 44.3).

### Inter-chromosomal phasing

#### Genome tiling by IBD segments

Our inter-chromosomal phasing approach begins by ‘tiling’ the genome with IBD segments. We define a tile as a set of overlapping IBD segments (Methods) and we assume that tiles are well phased in the Beagle or SHAPEIT pre-phasing step. To confirm this, we computed PER*_intra_*for each tile from the UKB and 23andMe trio children (Figure 7). Here, we are not concerned about the phase of a focal individual’s tiles relative to each other, but only the phase quality within a single tile. We first note that PER*_intra_* is low for all non-Asian ancestry groups, ranging from means of 0.10% in admixed African-ancestry 23andMe individuals to 0.25% in the UKB to 0.41% in Ashkenazi 23andMe individuals. These mean PER*_intra_* values are actually lower than their respective chromosome 2 SER when including blips (PER*_intra_*includes blips), except for the Ashkenazi. The low PER*_intra_* values we observe in all non-Asian ancestry groups indicate one of two things about the switch errors in them: (1) these switch errors occur close to the ends of tiles and/or (2) there are pairs of switch errors separated by one (a blip) or a few heterozygous sites. In either scenario, the phased haplotype is largely uninterrupted. In fact, the median PER*_intra_* for all non-East Asian trio children is zero (meaning the haplotypes match the transmitted parental haplotypes exactly). Mean PER*_intra_*values in 23andMe East and South Asians are much higher at 2.87% and 3.77%, respectively. This suggests that the haplotypes derived from IBD tiling are still mostly intact, but motivates further inquiry. Overall, these results suggest that IBD is powerful for identifying chromosomal regions that are well-phased, particularly in populations that are well-represented in the pre-phasing step. We also note a pattern that longer tiles have lower PER*_intra_* (Figure 7), although there could be a number of reasons for this. For example, the ends of tiles correspond to the ends of IBD segments (which are likely inexact) meaning there may be a few sites on the ends of the tiles that are not phased correctly; these are a smaller percentage of long tiles. Additionally, non-Asian 23andMe trio tiles are, on average, much longer than the UKB tiles: European 23andMe tiles have mean 918 heterozygous SNPs (sd 939 SNPs) versus 513 SNPs (sd 427) for UKB tiles. 23andMe South Asian tiles are the smallest (mean 325 SNPs; sd 355 SNPs).

**Figure 7.**
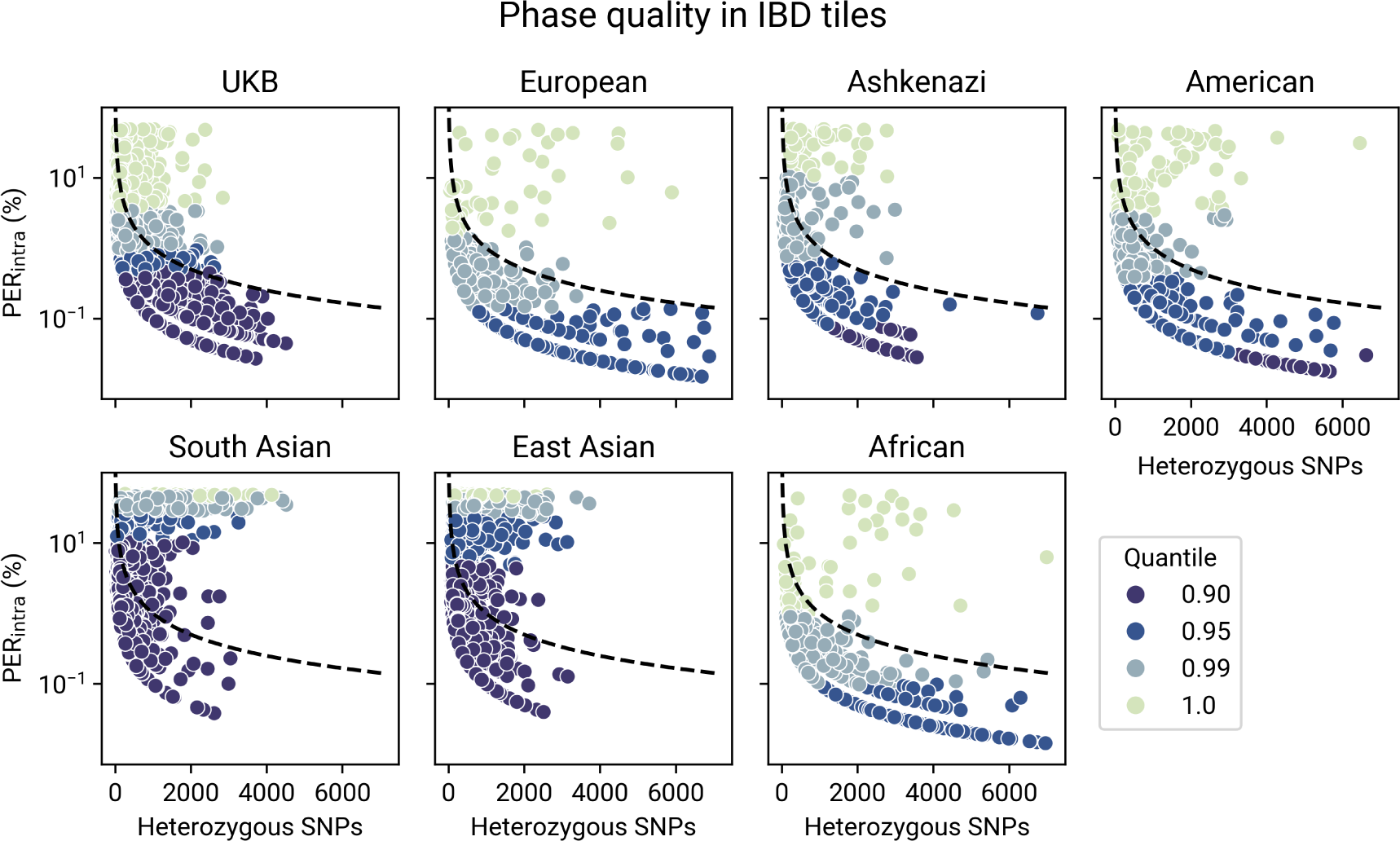
The phase quality in IBD tiles, as measured by PERintra. Each point represents a tile, plotted according to its length (the number of heterozygous sites in the tile; x-axis) and PERintra (y-axis). The dashed line represents PERintra when there are 10 mismatches between a tile’s haplotype and the true haplotype. Each point is also colored by the quantile the tile’s PERintra falls in (computed by ancestry group).

After tiling the genome, we create a graph *G* which is a set of nodes (IBD segments) that are either in the same tile or correspond to the same relative (Methods). In practice, *G* may be a collection of connected components; these correspond to IBD from a disjoint sets of relatives in which, at least in our scheme, there is no information about how the tiles should be phased relative to each other. We perform spectral clustering on *G*_1_, which is the largest connected component in terms of the number of nodes. The average UK Biobank trio has *G* covering 3404 cM of the genome (std. dev 287.85 cM), which corresponds to 90.9% of the genome. However, *G*_1_ covers an average of 2870 cM (std. dev 504.96 cM), which is 76.6% of the genome. In the 23andMe trios, *G*_1_ covers, on average, more of the genome (see Table S5): 84.5% for Europeans (sd 15.92%), 82.2% for admixed American (sd 16.12%), and 85.9% for admixed African-ancestry (sd 11.68%). The numbers are much lower for the 23andMe Asian groups: 27.1% for East Asian (sd 24.9%) and 29.3% for South Asians (sd 19.2%).

#### Evaluating inter-chromosomal phasing

We have shown that tiles represent regions of the genome that are well-phased, and that tiles cover significant portions of the genome for most non-Asian ancestry trio children. Our next task to perform inter-chromosomal phasing is to phase the tiles relative towards to each other. The important step in constructing *G* is in deciding whether to add positively-weighted edges between IBD segments that are in different tiles and belong to the same relative. Adding a positively-weighted edge in such a case indicates a belief that the IBD segments were inherited from the same parent and, because we know which haplotype index the IBD segments fall on, allows us to either keep the default phase of the two tiles or flip the phase of one of the tiles. We introduced two thresholds that determine if a positive edge should be added between tiles: *t*_1_ (both segments must be longer than this threshold) and *t*_2_ (the relative must share a total of *t*_2_ cM IBD with the focal individual). Increasing these thresholds is more conservative, adding fewer edges and connecting fewer tiles, and so represents a trade-off between accuracy and call rate (the percentage of sites that we are able to phase). This is clear in the UK Biobank British trio children, where PER*_inter_* steadily climbs with call rate (Figure S13). We ran the method on all combinations of *t*_1_ *∈ {*5, 10} and *t*_2_ *∈ {*10, 25, 50, 75, 100}, except for the degenerate combination (*t*_1_*, t*_2_) = (10, 10). We find that the higher, more conservative thresholds do phase with lower error rates, but phase fewer sites, e.g., compare Table 1 and Table S5.

**Table 1.** HAPTIC’s PER*_inter_* for different ancestry groups for heterozygous sites in IBD tiles. Shown are the results for the thresholds (*t*_1_ and *t*_2_) that result in the lowest PER*_inter_*. We also show the call rate, which is the percentage of heterozygous sites phased.

| Ancestry group | $t_1$ | $t_2$ | $PER_{inter}$ (%) | | | call rate (%) | | |
| --- | --- | --- | --- | --- | --- | --- | --- | --- |
|  |  |  | mean | std | median | mean | std | median |
| UKB British | 10 | 100 | 0.55 | 3.1337 | 0.0841 | 33.49 | 24.02 | 29.56 |
| South Asian | 5 | 100 | 29.28 | 16.65 | 35.57 | 19.66 | 14.09 | 16.92 |
| admixed American | 10 | 100 | 5.76 | 12.46 | 0.02 | 65.64 | 23.31 | 69.14 |
| European | 10 | 75 | 2.43 | 6.31 | 0.02 | 67.13 | 22.87 | 74.37 |
| East Asian | 10 | 50 | 21.86 | 18.41 | 18.85 | 23.49 | 21.02 | 14.48 |
| Ashkenazi | 10 | 100 | 31.65 | 11.91 | 31.62 | 51.43 | 7.23 | 51.19 |
| admixed African | 10 | 100 | 2.83 | 6.85 | 0.015 | 59.43 | 23.68 | 64.45 |

Figure S16 shows the distribution of PER*_inter_*values for each combination of thresholds. Table 1 shows the lowest PER*_inter_*achieved for each ancestry group: the lowest mean PER*_inter_* achieved is 0.55% (median 0.08%; sd 3.1%) for UKB trio children at thresholds *t* = (*t*_1_*, t*_2_) = (10, 100), which phases 33.5% of sites (sd 24%). However, given the trade-off between call rate and PER*_inter_*, it is difficult to decide the “best” threshold; e.g., a better threshold might be *t* = (*t*_1_*, t*_2_) = (5, 50), which has a mean PER*_inter_* of 1.72% (median 0.13%, sd 5.44%) but phases 44% of sites (sd 24.8%). Overall, the distribution of PER*_inter_* has a large tail: most individuals are very well-phased, but there are some that are very poorly phased ([29] also observe this in their inter-chromosomal phasing method). For 23andMe European ancestry individuals, for example, a mean PER*_inter_* of 2.43% at *t* = (*t*_1_*, t*_2_) = (10, 75) (mean call rate of 67%) is achieved, but the median PER*_inter_* is 0.02% (median call rate is 74%). PER*_inter_* and call rates are similar for 23andMe admixed American and admixed African ancestry groups (median PER*_inter_* 0.02% and 0.015%, respectively, and median call rate 69.14% and 64.45%, respectively). Both the mean and median PER*_inter_*are high for the Asian groups: median 18.9% for East Asians and 35.6% for South Asians. The performance in the 23andMe Ashkenazi individuals is also poor: despite the tiles being well-phased and tiles covering most of the genome, PER*_inter_* is high with a median of 31.64%. Also recall that in our intra-chromosomal phasing analysis that Ashkenazi trio children were among the ancestry groups with the lowest SER on chromosome 2 (Table S2). The poor performance is likely due to the presence of necessarily close, non-descendant relatives who are related to both parents.

The PER*_inter_* we have reported is for regions covered by IBD tiles only. Figure S15 reports PER*_inter_* for all chromosomes covered by IBD tiles. We perform a rudimentary phasing algorithm for regions on the chromosome that are not covered by an IBD tile. If the region is at the beginning of the chromosome, it takes the phase of the downstream tile (e.g., if the downstream tile’s phase needs to be switched, we switch the phase of the region). If the region is between two tiles (or between a tile and the end of the chromosome), we switch the phase if the upstream tile’s phase needs to be switched. While this is not statistically motivated (e.g., SHAPEIT scaffold phasing would be preferred), it is fast and works well when whole chromosomes are well-phased, which may be the case in many 23andMe samples. In UKB, for example, this does not work well as its median PER*_inter_* is 13.1% (median call rate 93.3%). However, this works remarkably well in 23andMe where the median PER*_inter_* is 0.92% (median call rate 93.8%) for European-ancestry individuals and 0.09% in the admixed African-ancestry individuals (median call rate 92.7%). We also show PER*_inter_* for all sites in the genome, including sites on chromosomes that have no tiles, in Figure S14. These error rates are much higher, particularly for individuals with few tiles, as chromosomes without tiles are phased at random with respect to the tile phase.

#### Detecting descendants

We outlined a procedure for detecting descendants, which involves applying our inter-chromosomal phasing method on sets of relatives that exclude potential descendants (**Methods**). We expect our ability to detect descendants to improve when we can inter-chromosomally phase more of the genome without these potential descendants, but this may be difficult because potential descendants are close relatives that are highly informative for this phasing. Overall, we are able to detect 86.8% of simulated UKB descendants. However, for focal individuals in which we are able to phase > 200 cM of the genome with potential descendants excluded, this increases to 94.0% (and to 99.0% when 1,000 cM of the genome is phased). A false positive descendant is a close non-descendant relative who is classified as a descendant; we found a false positive rate of only 0.54%.

## DISCUSSION

Accurate haplotype phasing is crucial for a wide range of genetic analyses, and our findings highlight both the progress and remaining challenges in achieving high-quality phasing across diverse populations. In the first part of our paper, we benchmarked two popular state-of-the-art phasing methods, SHAPEIT and Beagle, on ∼6,000 diverse trio children using a reference panel of 8.6M research-consented 23andMe customers, representing the largest published benchmarking of the two methods. Mean SER with blips removed in non-Asian trio children are extraordinarily low, representing only a handful (< 6) switch errors. Further, we find that 56.8% of SHAPEIT-phased non-Asian 23andMe trio children have *zero* non-blip switch errors on chromosome 2 (Figure S7), meaning that nearly the entirety of the chromosome is phased perfectly, except for consecutive switch errors which do not alter the phase of the surrounding haplotypes and may be caused by genotype error [3]. We ran the same analysis with 1,606 trio children from a merged UK Biobank-TGP dataset, using the merged dataset (with trio parents removed) as a reference panel. The 985 British-ancestry trio children were phased at a comparable SER to 23andMe European-ancestry trio children when blips were removed, but performance in non-British trio children was generally poor, with some samples having 1500+ non-blip switch errors. With the merged UKB-TGP dataset, we also performed analyses demonstrating that IBD sharing reduces SER (Figure S10) and finding an association between switch errors and regions with low IBD coverage (Figure S11).

We also developed a new method for inter-chromosomal phasing, HAPTIC. The goal of our method is to identify the alleles on each parental haplotype for all chromosomes in a focal individual. HAPTIC takes as input IBD segments that were called using genotype data pre-phased by either SHAPEIT or Beagle. Our results from the first half of the paper suggest that short-range haplotype phase quality is high in large datasets, particularly for individuals that share IBD with many others in the dataset. We take advantage of this high phase quality, and use overlapping IBD segments to identify haplotypes that have been well-phased in the pre-phasing step. We show that, indeed, these haplotypes, which we call IBD tiles, are well-phased (Figure 7). This scheme reduces the task of phasing single sites into phasing the IBD tiles relative to each other, which we do using a spectral clustering technique introduced by [24]. We use a simple statistic called the inter-chromosomal phase error rate (PER*_inter_*) to assess our method. PER*_inter_*measures the Hamming distance between the variants in the inferred parental haplotype set against the variants in the true (transmitted) parental haplotype set. PER*_inter_* is much harsher than SER, which only compares the phase of consecutive heterozygous sites (Figure 1). In regions covered by IBD tiles, HAPTIC achieves low PER*_inter_* in several 23andMe groups (European, African, admixed American) and in the white British UKB trio children; in fact, median PER*_inter_*in tiled regions is actually lower than median chromosome 2 SER. However, HAPTIC results in more variable performance than intra-chromosomal phasing methods. HAPTIC performs poorly in South Asian and East Asian trio children, likely because these groups have overall lower sample sizes and consequently less IBD in the 23andMe dataset. Despite high-quality intra-chromosomal phasing and lots of IBD, HAPTIC performs poorly in Ashkenazi Jewish trio children. Given that our method filters out IBD segments in regions of IBD2 (which filters long regions of homozygosity), the poor performance is not due to regions of homozygosity that result from consanguinity (it is well-documented that consanguinity levels in Ashkenazi Jewish populations are high) [21]. More likely, the poor performance is due to pedigree collapse, which may be associated with consanguinity deeper in the genealogy.

Overall, our study highlights the importance of IBD in haplotype phasing, whether used implicitly (Beagle and SHAPEIT) or explicitly (HAPTIC). IBD coverage, particularly coverage of many long IBD segments (that are typical of close relative IBD sharing), is vital for HAPTIC, which uses the IBD segments explicitly. The negative correlation we find between IBD coverage and SER is striking, but we caution that our IBD coverage statistic does not include the many IBD segments present that are < 3cM, which would still contribute to (and lower) short-range phase quality statistics such as SER. Importantly, IBD coverage can act as a measure of representation in the dataset in a way that is more useful (e.g., for predicting phase accuracy) than the information contained in continental labels. For instance, we find that SER for UKB self-described Indians (which have high IBD coverage) is much lower than TGP Telegu Indians from the UK (which have low IBD coverage), despite both groups being described as ‘Indian’. Telegu speakers in the UK number approximately 31,000 (whereas more common South Asian languages are spoken among hundreds of thousands in the UK) [12], so Telugu’s may not be well-represented in the UK Biobank. Using IBD coverage to assess representation also eliminates the need for ancestry labels, which are difficult to define.

A formidable challenge we encountered in developing HAPTIC is in trying to balance accuracy and call rate (the percentage of heterozygous sites genome-wide phased). HAPTIC requires relatives that share multiple IBD segments with a focal individual in order to connect haplotypes across chromosomes. In the UKB, most of these multi-segment relatives are distantly related (e.g., sharing two 5 cM IBD segments with the focal individual). We found that, for 35% of these distant relatives (not shown), the two IBD segments were inherited through different parents, so distant relatives could not be used to reliably connect haplotypes across the genome. HAPTIC’s spectral clustering technique clusters a signed graph of IBD segments into sets of paternally- and maternally-inherited segments. A key step in HAPTIC is assigning edge weights between segments corresponding to the same relative. The edge weight should reflect the certainty that the IBD segments are inherited through the same parent. For example, a pair of long IBD segments shared with the same relative are likely inherited through the same parent. We use two thresholds to determine whether to add a positive edge-weight: *t*_1_ (both IBD segments must be longer than *t*_1_) and *t*_2_ (the total kinship of the relative to the focal must be larger than *t*_2_). In order to achieve high accuracy, the thresholds must be set high, but this indiscriminately removes all edges between IBD segments of more distant relatives, even if some of the edges would be useful for clustering the haplotypes. Thus, our method does not achieve a high call rate in the UK Biobank, but we note that the regions in between the IBD tiles (or between chromosome ends and IBD tiles) can be statistically-phased with SHAPEIT, using the phased haplotypes as a scaffold, as done in [20]. Another limitation is that our method does not infer the sex of the parent that transmitted the haplotype sets. This can be done with relative ease in male focal individuals by looking at IBD on the X-chromosome [20], i.e., a male’s IBD tiles on the X-chromosome are maternal haplotypes. Otherwise, without external genealogical information (that could label relatives and their associated IBD as maternal or paternal), inferring the parent sexes is non-trivial; one potential avenue would be in analyzing the breakpoints of the IBD tiles and associating them with the male and female sex-specific recombination map, similar to the work of [31] which used sex-specific recombination maps to distinguish maternal and paternal half-siblings.

Hofmeister et al. (2021) [20] already showed a use-case for inter-chromosomal phasing for performing parent-of-origin effect GWA studies. Another potential use-case of inter-chromosomal phasing is in parent-of-origin specific local ancestry maps. For instance, the output of our method can easily be merged with local ancestry inference (LAI) data. This could potentially be used to refine the timings of migration [15]. While current applications of inter-chromosomal phasing are still emerging, the increasing availability of large and diverse datasets, coupled with advancements in phasing methodologies like HAPTIC, will yield new insight into a range of population genetic and statistical genetic problems.

## DATA AND CODE AVAILABILITY

HAPTIC will be made available upon publication.

## ACKNOWLEDGEMENTS

We would like to thank the research participants and employees of 23andMe for making this work possible.

The following members of the 23andMe Research Team contributed to this study: Stella Aslibekyan, Adam Auton, Elizabeth Babalola, Robert K. Bell, Jessica Bielenberg, Ninad S. Chaudhary, Zayn Cochinwala, Sayantan Das, Emily DelloRusso, Payam Dibaeinia, Sarah L. Elson, Nicholas Eriksson, Chris Eijsbouts, Teresa Filshtein, Pierre Fontanillas, Davide Foletti, Will Freyman, Zach Fuller, Julie M. Granka, Chris German, Éadaoin Harney, Alejandro Hernandez, Barry Hicks, David A. Hinds, M. Reza Jabalameli, Ethan M. Jewett, Yunxuan Jiang, Sotiris Karagounis, Lucy Kaufmann, Matt Kmiecik, Katelyn Kukar, Alan Kwong, Keng-Han Lin, Yanyu Liang, Bianca A. Llamas, Aly Khan, Steven J. Micheletti, Matthew H. McIntyre, Meghan E. Moreno, Priyanka Nandakumar, Dominique T. Nguyen, Jared O’Connell, Steve Pitts, G. David Poznik, Alexandra Reynoso, Shubham Saini, Morgan Schumacher, Leah Selcer, Anjali J. Shastri, Jingchunzi Shi, Suyash Shringarpure, Keaton Stagaman, Teague Sterling, Qiaojuan Jane Su, Joyce Y. Tung, Susana A. Tat, Vinh Tran, Xin Wang, Wei Wang, Catherine H. Weldon, Amy L. Williams, Peter Wilton.

All UKB-TGP analyses were conducted using computational resources and services at the Center for Computation and Visualization, Brown University.

## SUPPLEMENTARY MATERIALS

**Figure S1.**
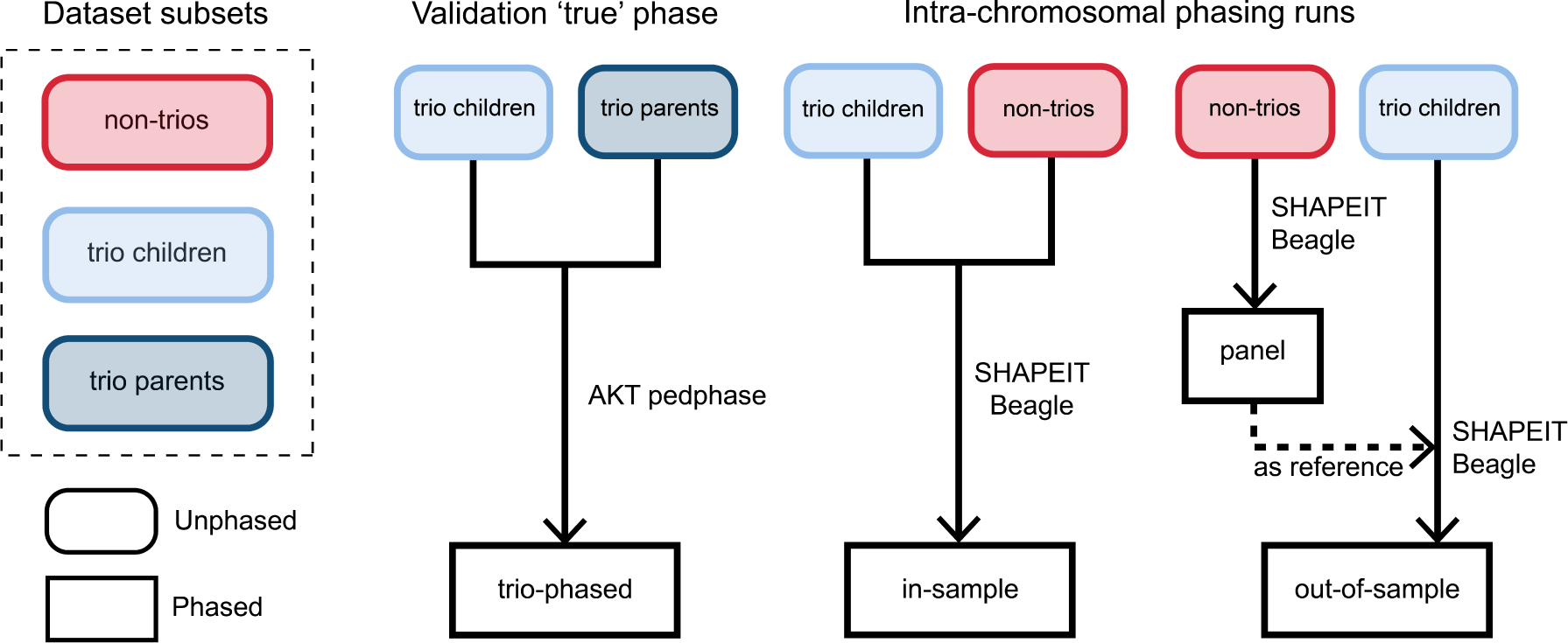
Intra-chromosomal phasing scheme. We split the full dataset into three subsets: trio children (individuals with both parents in the dataset), trio parents (the parents of trio children), and non-trios (all other samples). The graphic describes which data subsets are used in three different phasing runs: the validation trio-phasing run and two different statistical phasing runs (‘in-sample’ and ‘out-of-sample’ phasing).

**Figure S2.**
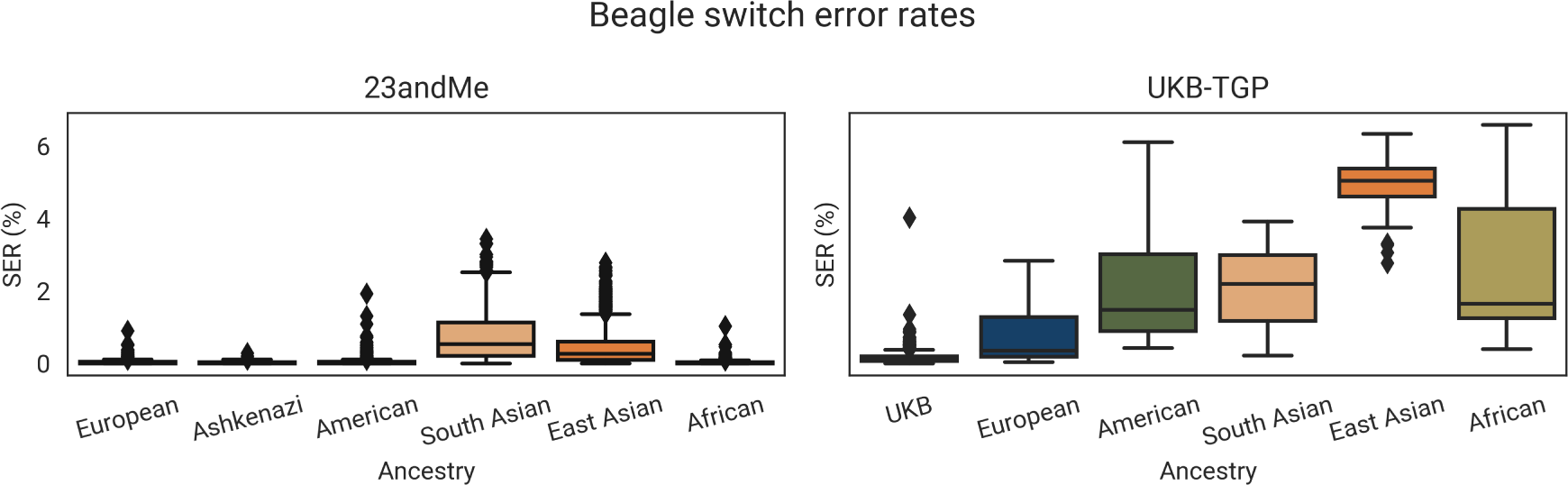
Beagle switch error rates (SER) including phasing ‘blips’ for 23andMe trio children (left) and UKB-TGP trio children (right), split by ancestry group.

**Figure S3.**
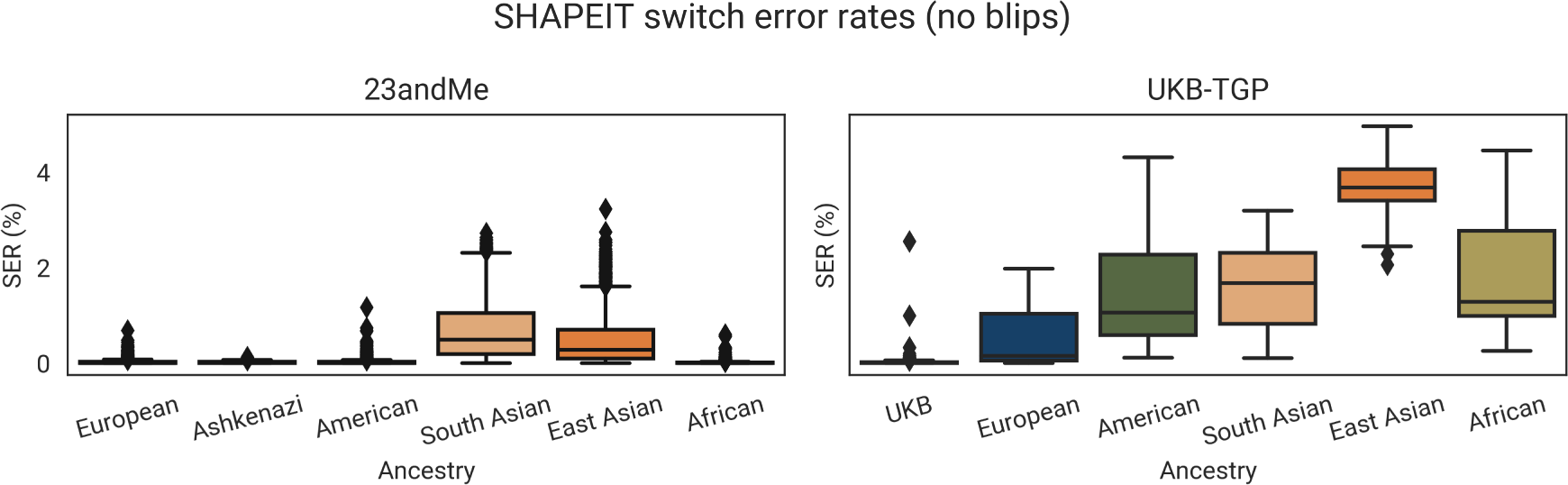
SHAPEIT switch error rates (SER) excluding phasing ‘blips’ for 23andMe trio children (left) and UKB-TGP trio children (right), split by ancestry group.

**Figure S4.**
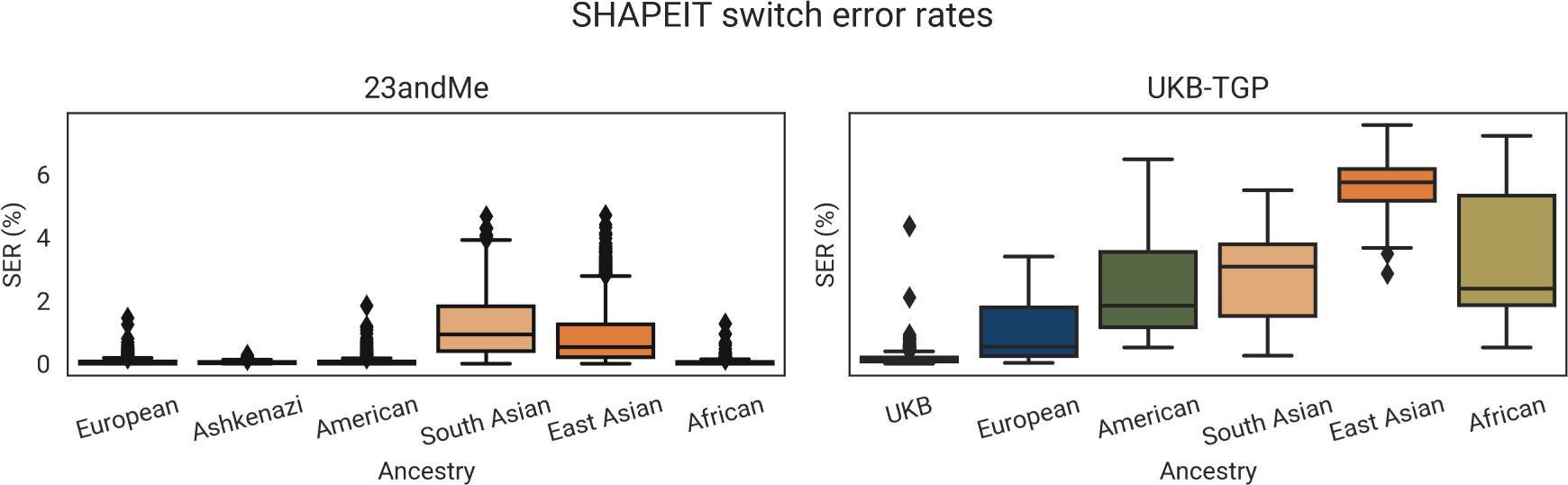
SHAPEIT switch error rates (SER) including phasing ‘blips’ for 23andMe trio children (left) and UKB-TGP trio children (right), split by ancestry group.

**Figure S5.**
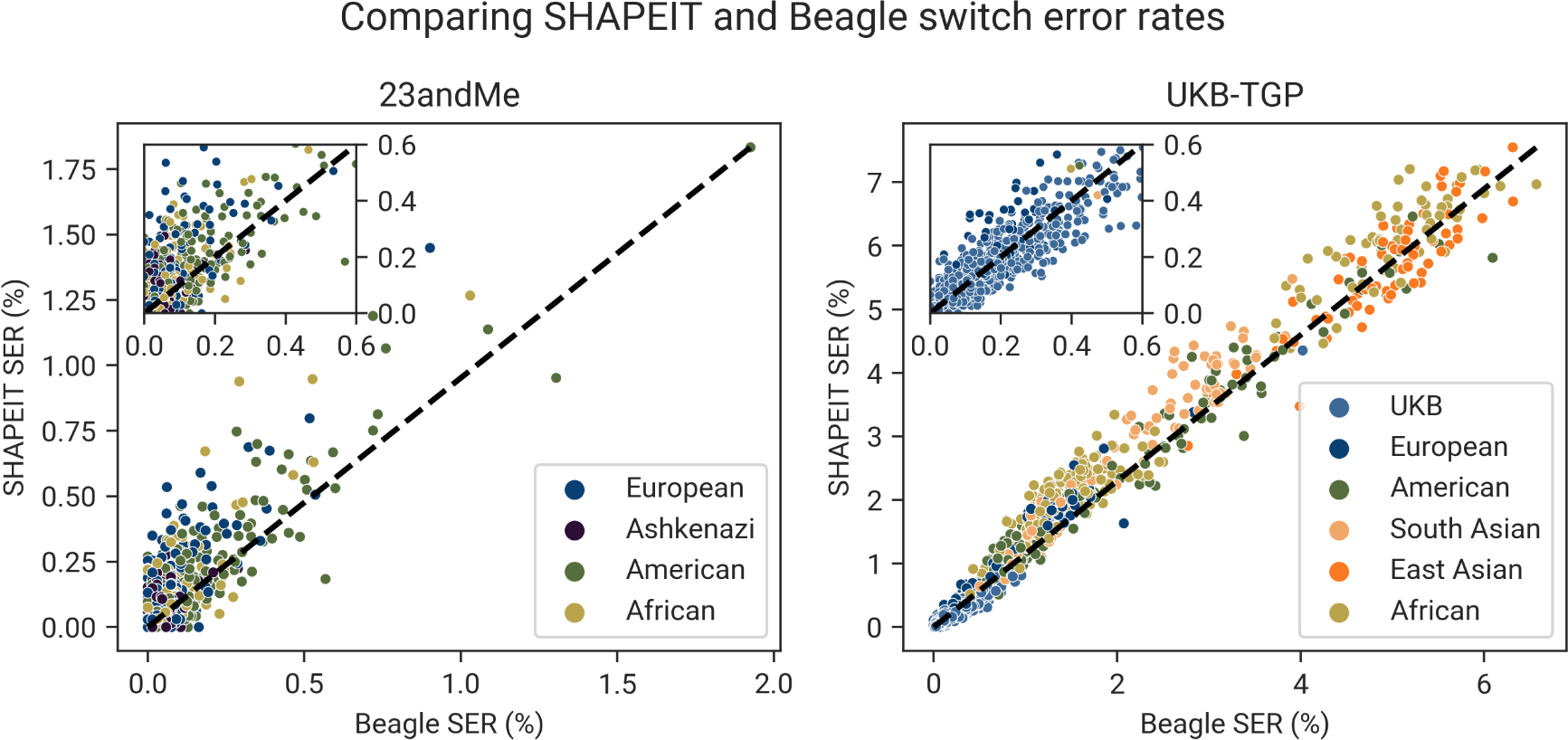
Comparison of Beagle switch error rates (SER; on x-axis) and SHAPEIT SER (y-axis) with phasing blips included. Each point is a trio child, colored by their ancestry group (note that this does not include South and East Asian trio children (see Figure S6 left panel). The dashed line indicates an equal Beagle SER and SHAPEIT SER. As most 23andMe trio children have SER < 0.60%, we included an inset showing these trio children.

**Figure S6.**
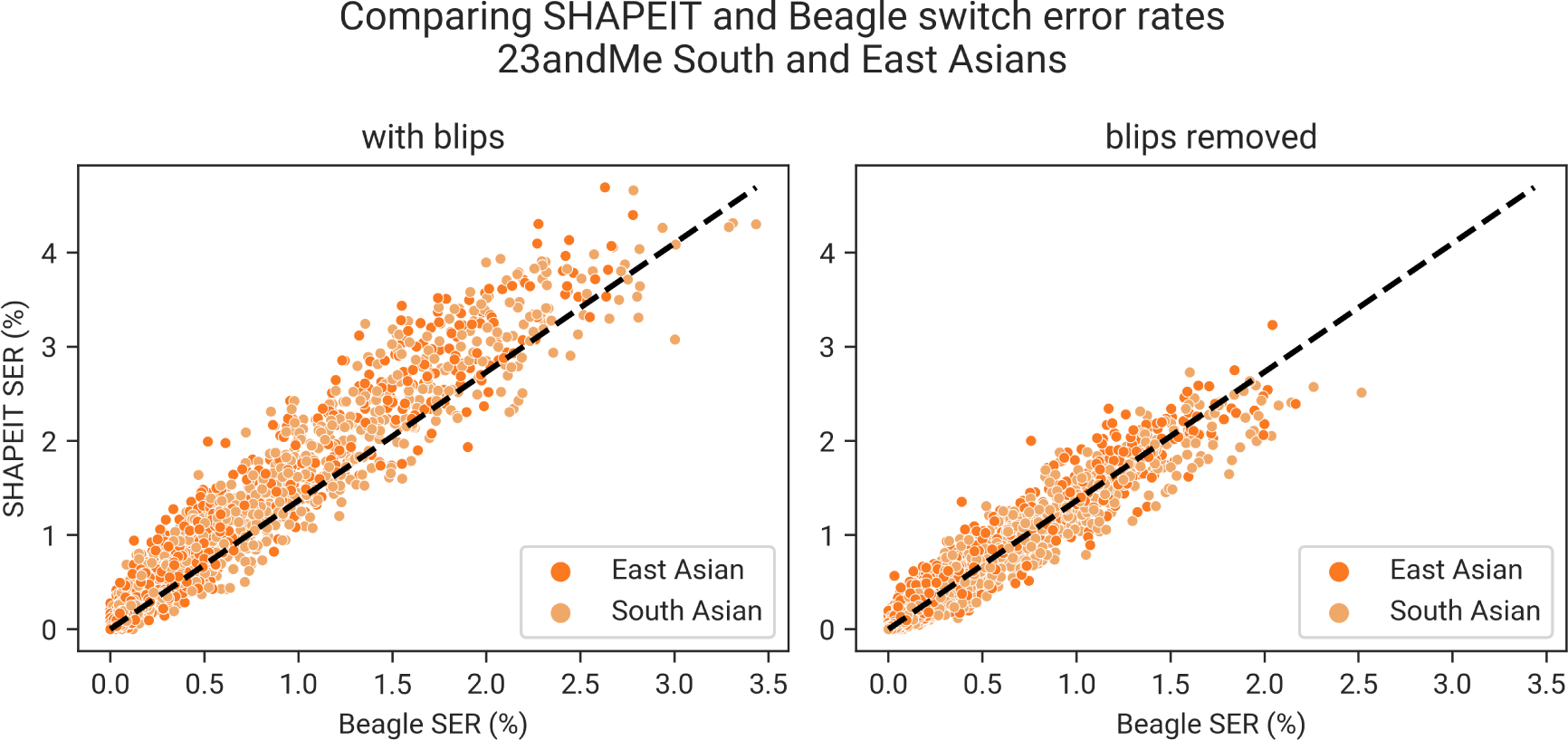
Comparison of Beagle switch error rates (SER; on x-axis) and SHAPEIT SER (y-axis) for South and East Asian ancestry trio children. The left panel SER includes phasing blips; the right panel SER is computed without phasing blips. The dashed line indicates an equal Beagle SER and SHAPEIT SER.

**Figure S7.**
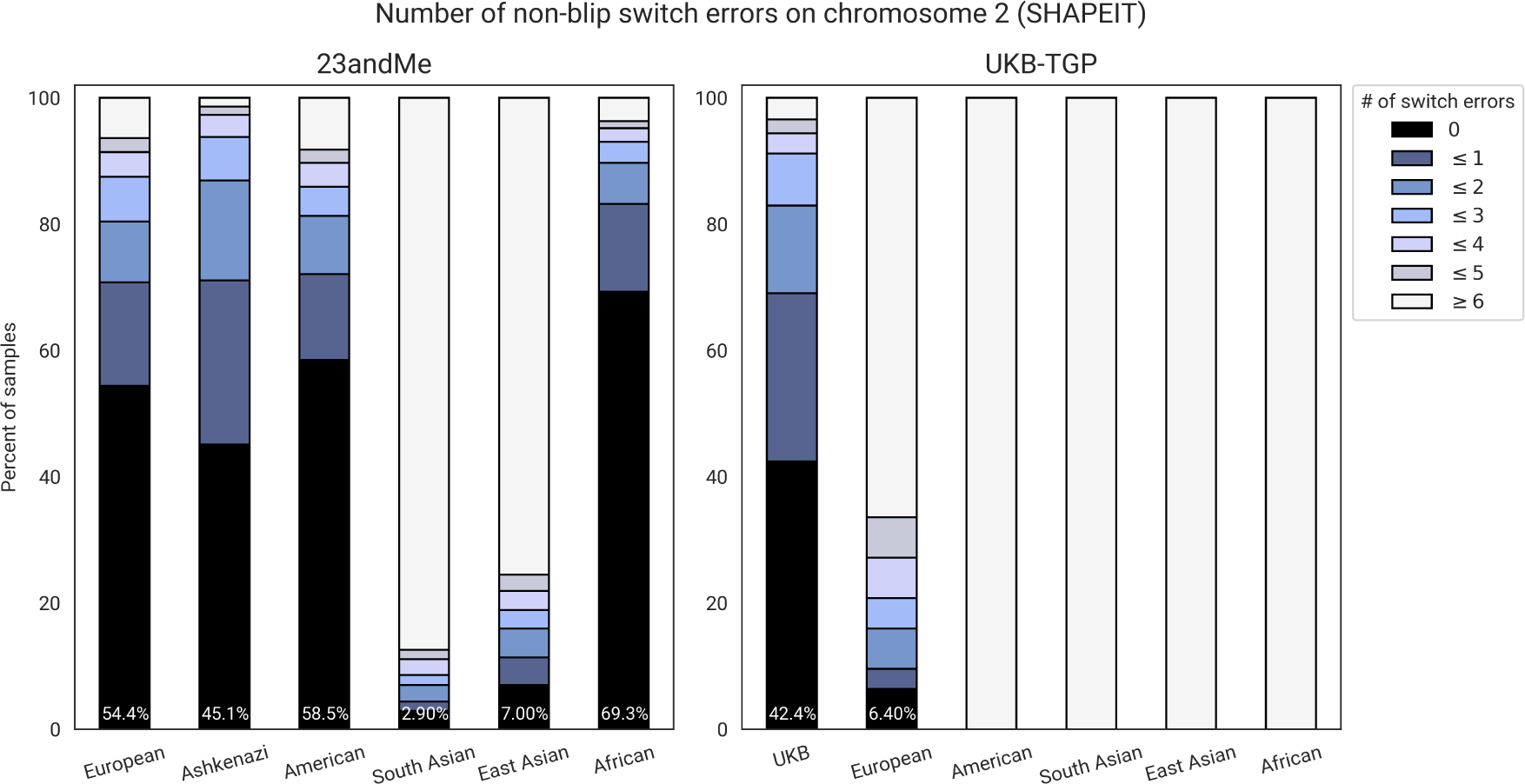
A stacked bar plot showing the distribution of the number of switch errors (SHAPEIT phasing; blips removed) in the trio children, split by dataset and ancestry group. For example, 54.4% of 23andMe European-ancestry trio children have zero non-blip switch errors. On the other hand, all admixed American UKB-TGP trio children have 6+ non-blip switch errors.

**Figure S8.**
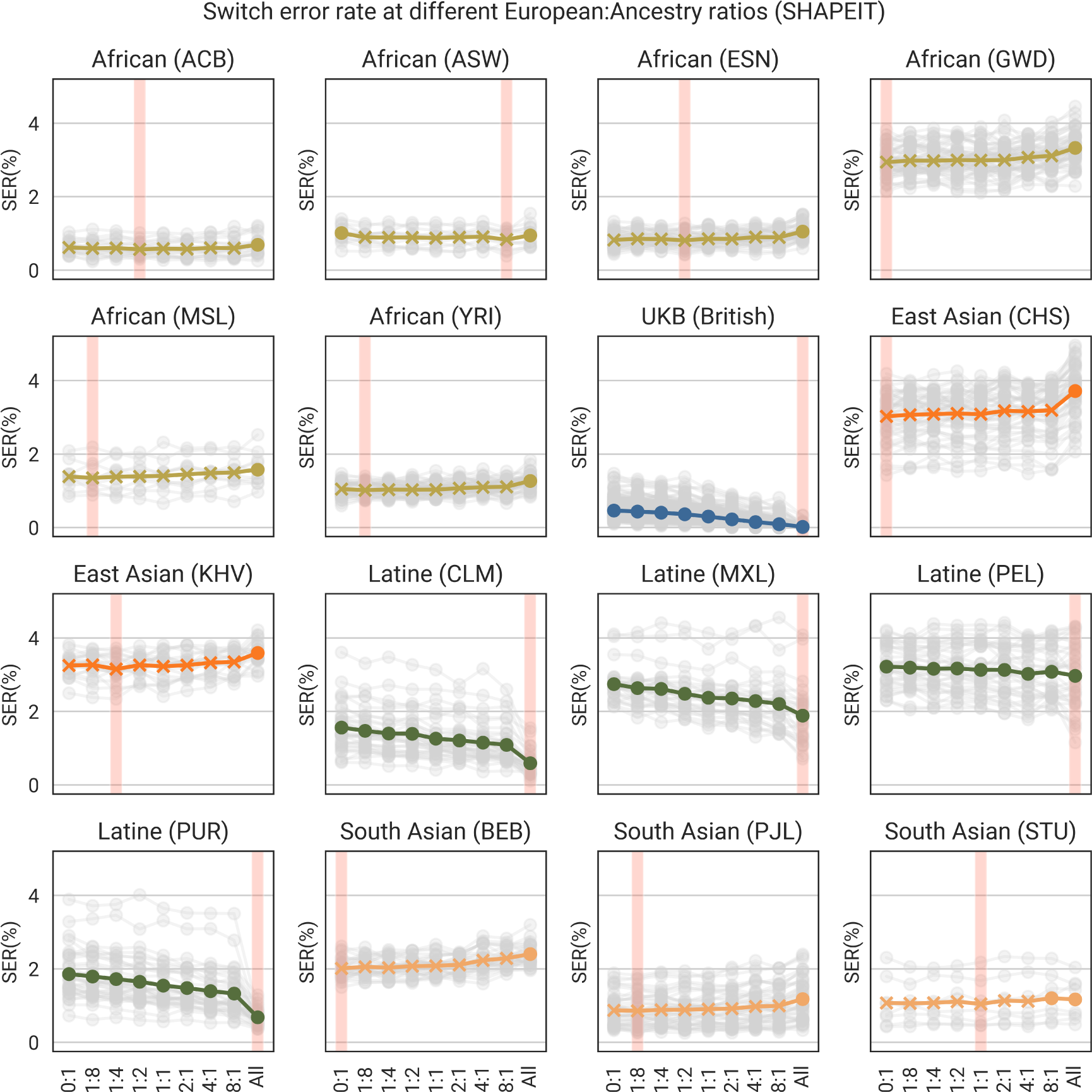
For each continental ancestry group, we did several SHAPEIT ‘in-sample’ phasing runs where we varied the number of Europeans in the dataset relative to the size of the ancestry group. For instance, the African 1:8 phasing run indicates that there was 1 European sample for every 8 African-ancestry samples. These ratios are on the x-axis (‘All’ is the in-sample run with the entire dataset). Here, we show the results split by 1000 Genomes Project population labels. Each trio child’s SERs are shown in grey, and the colored line is the mean line. ‘X’ markers indicate that the mean is below the ‘All’ (in-sample, entire dataset) mean. The red vertical bar indicates the ratio with the lowest mean SER for the population.

**Figure S9.**
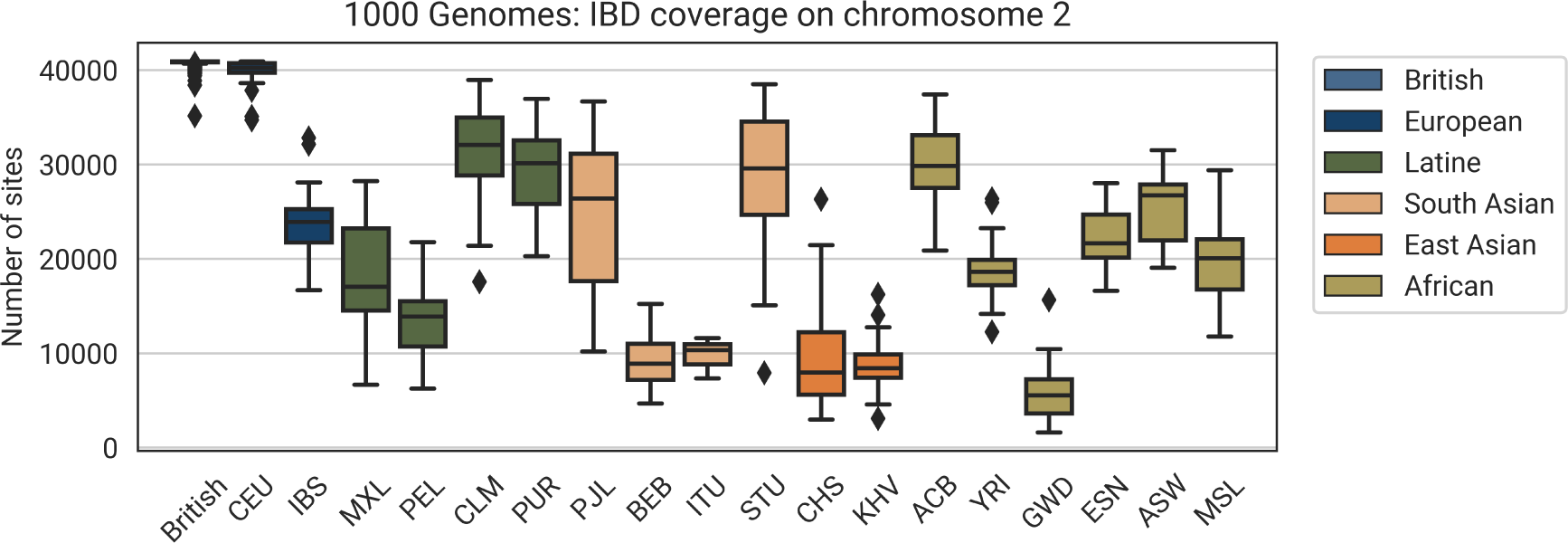
The distribution of IBD segment coverage, split by 1000 Genomes Project ‘Population’ label (except for ‘British’, which refers to UK Biobank British trio children). IBD coverage is the number of sites on chromosome 2 in which a trio child has at least one IBD segments covering.

**Figure S10.**
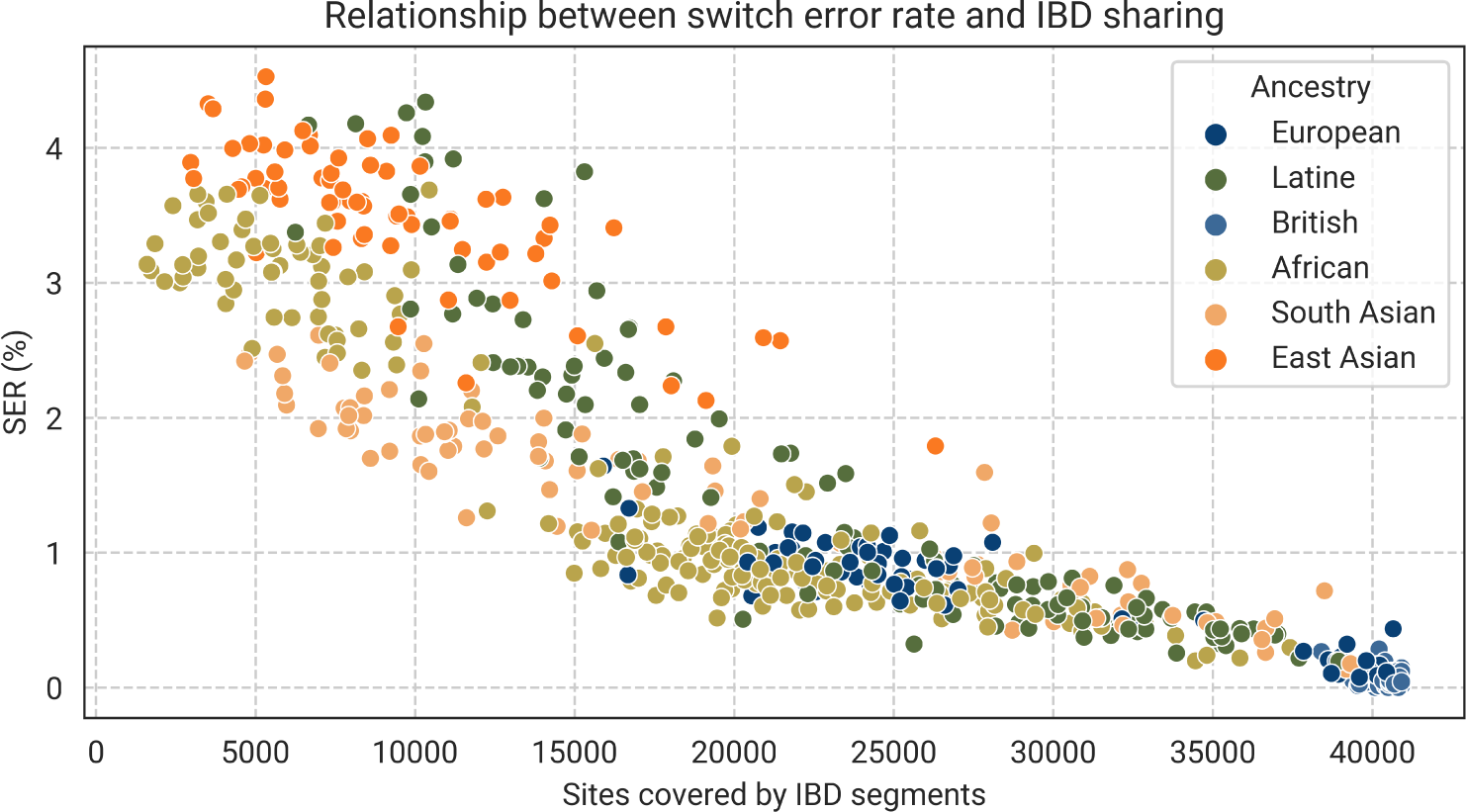
The relationship between switch error rate (SER) and IBD sharing. For each trio child, we computed the number of sites on chromosome 2 covered by at least one IBD segment and compare to the SER. We computed the IBD using trio-phased (Mendelian phasing) genotype data.

**Figure S11.**
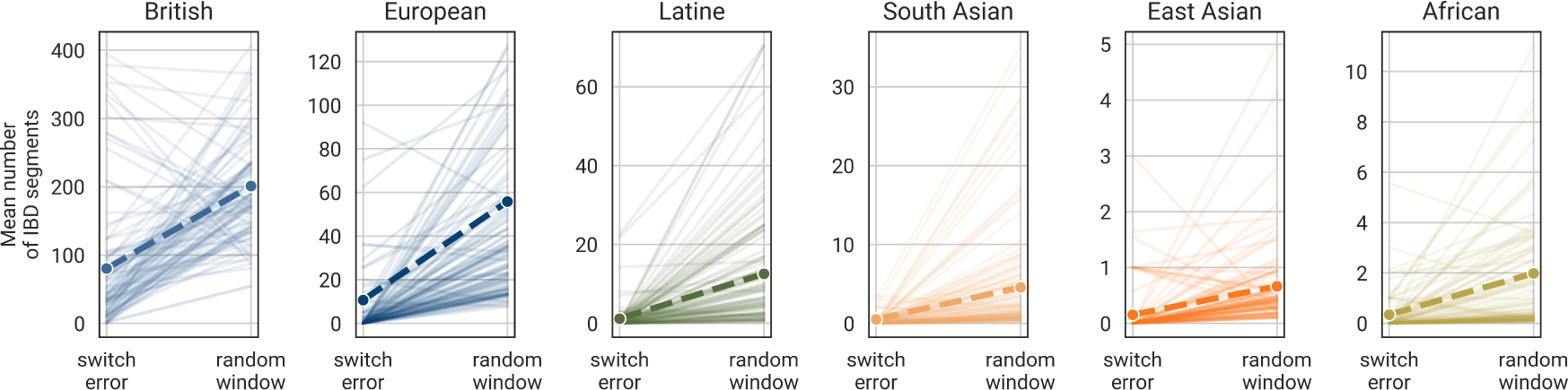
For each switch error for each individual, we computed the mean number of IBD segments that span the switch error. For each individual’s switch errors, we also computed the mean number of IBD segments that cover 100 random sites across the individual’s genome of the same span. Each line is one of 100 randomly selected switch errors, and the thick dashed line is the mean for all switch errors in the ancestry group. In general, we see the trend that switch errors occur in regions of lower IBD coverage than randomly selected regions.

**Figure S12.**
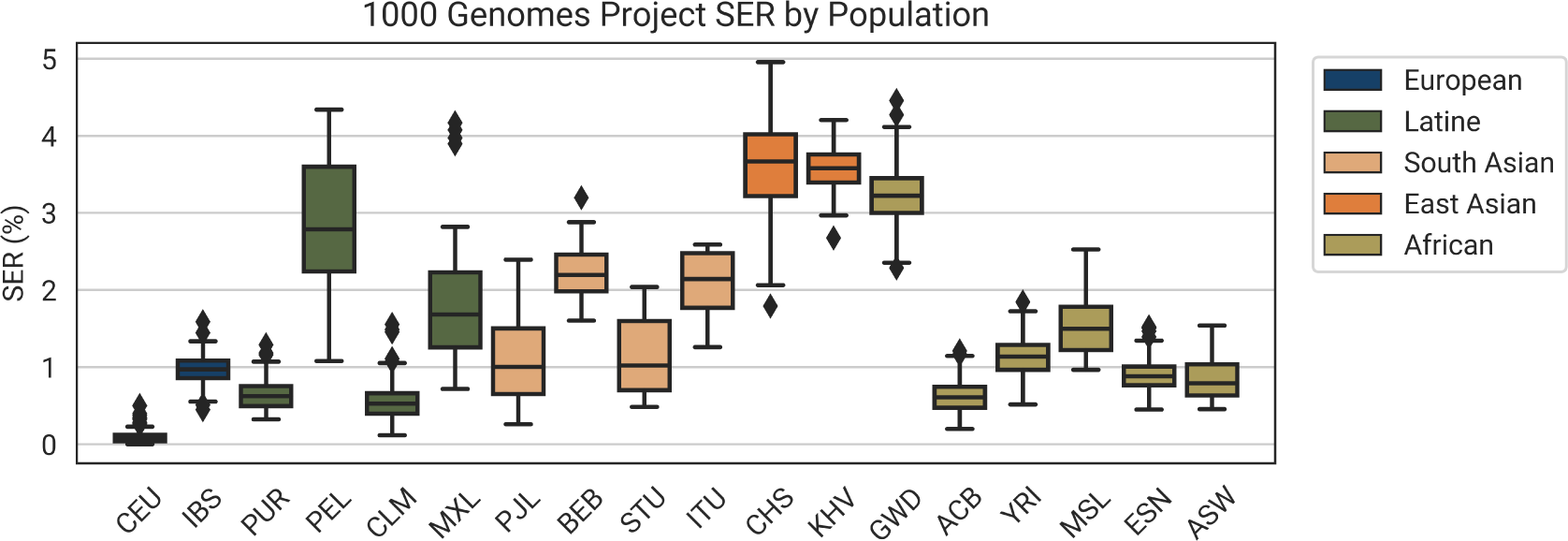
The Beagle switch error rate (SER) of 1000 Genomes Project populations. There is considerable variance in SER within the continental ancestry groups, which appears to be driven by representation in the UK Biobank.

**Figure S13.**
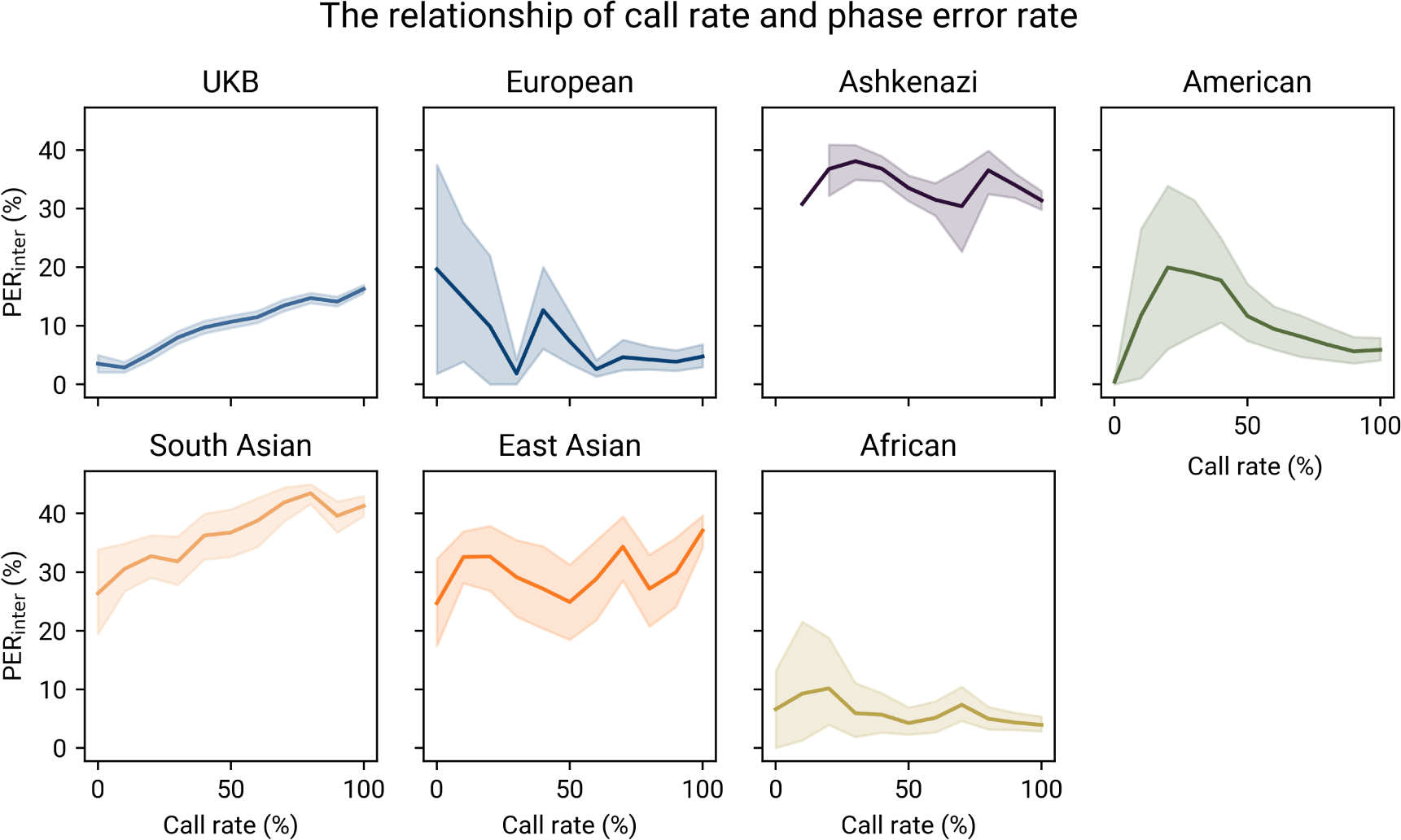
The relationship between the call rate (% of sites phased by our method) and the phasing accuracy, measured by PER*_inter_*and split by ancestry group. All non-UKB plots are 23andMe population plots. We binned each clustering run by call rate (bin widths 10%) and computed the mean PER*_inter_* in the bin. The 95% CI is shaded around the mean line.

**Figure S14.**
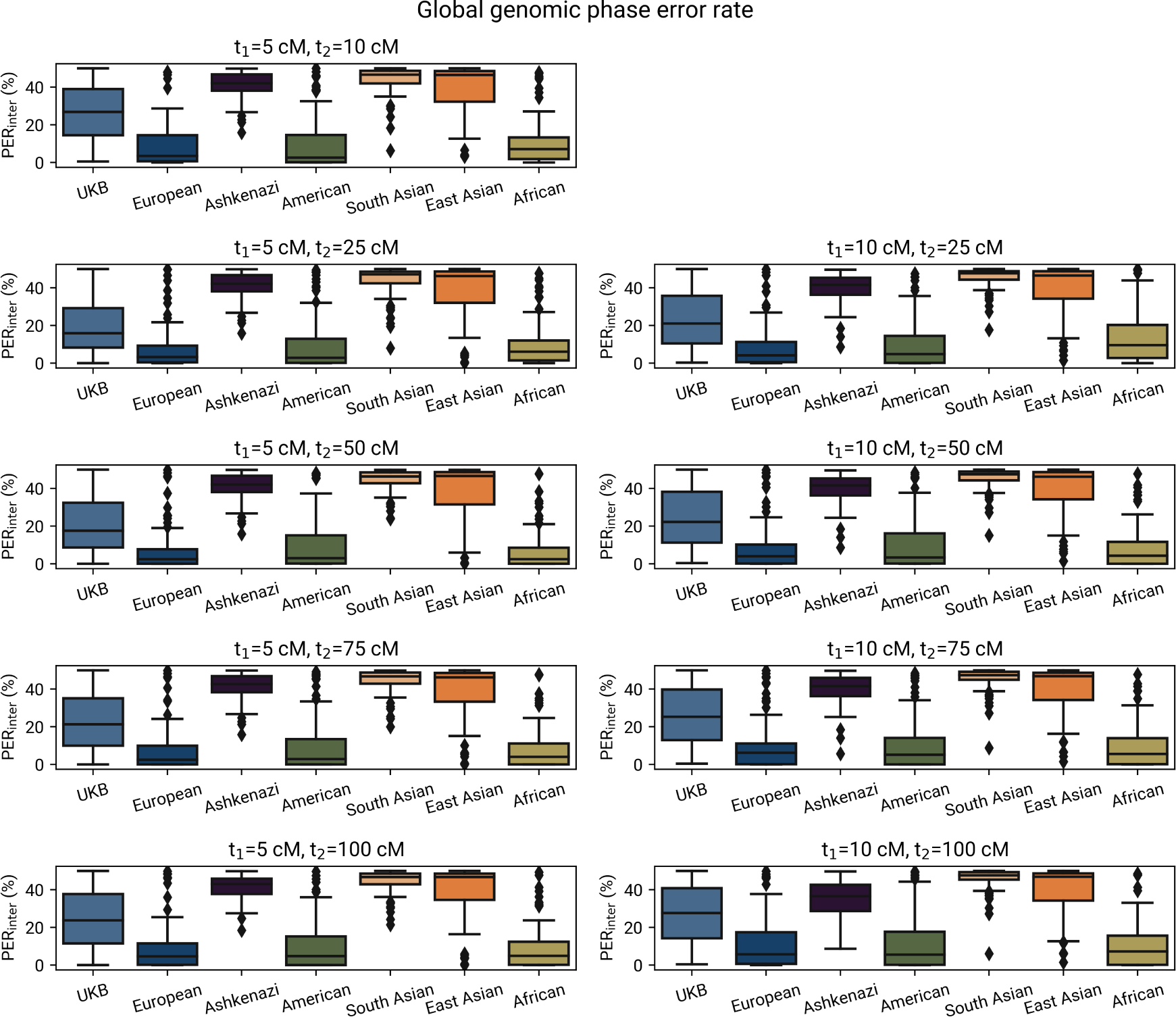
The phasing performance of HAPTIC as measured by PER*_inter_*, split by ancestry group (all non-UKB ancestry groups are 23andMe) for all sites in the genome, including those not covered by IBD tiles.

**Figure S15.**
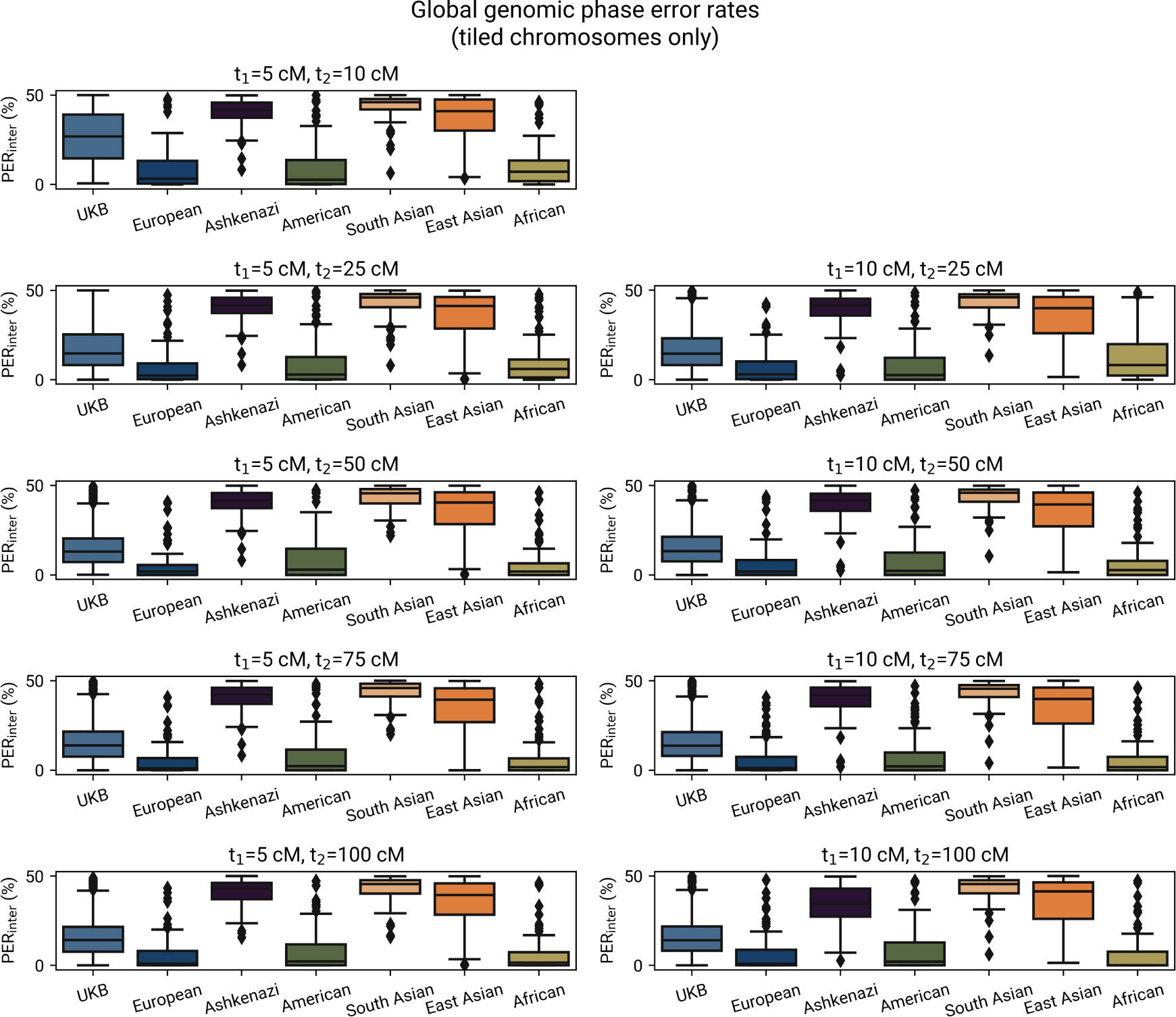
The phasing performance of HAPTIC in chromosomes covered by IBD tiles, as measured by PER*_inter_* and split by ancestry group (all non-UKB ancestry groups are 23andMe).

**Figure S16.**
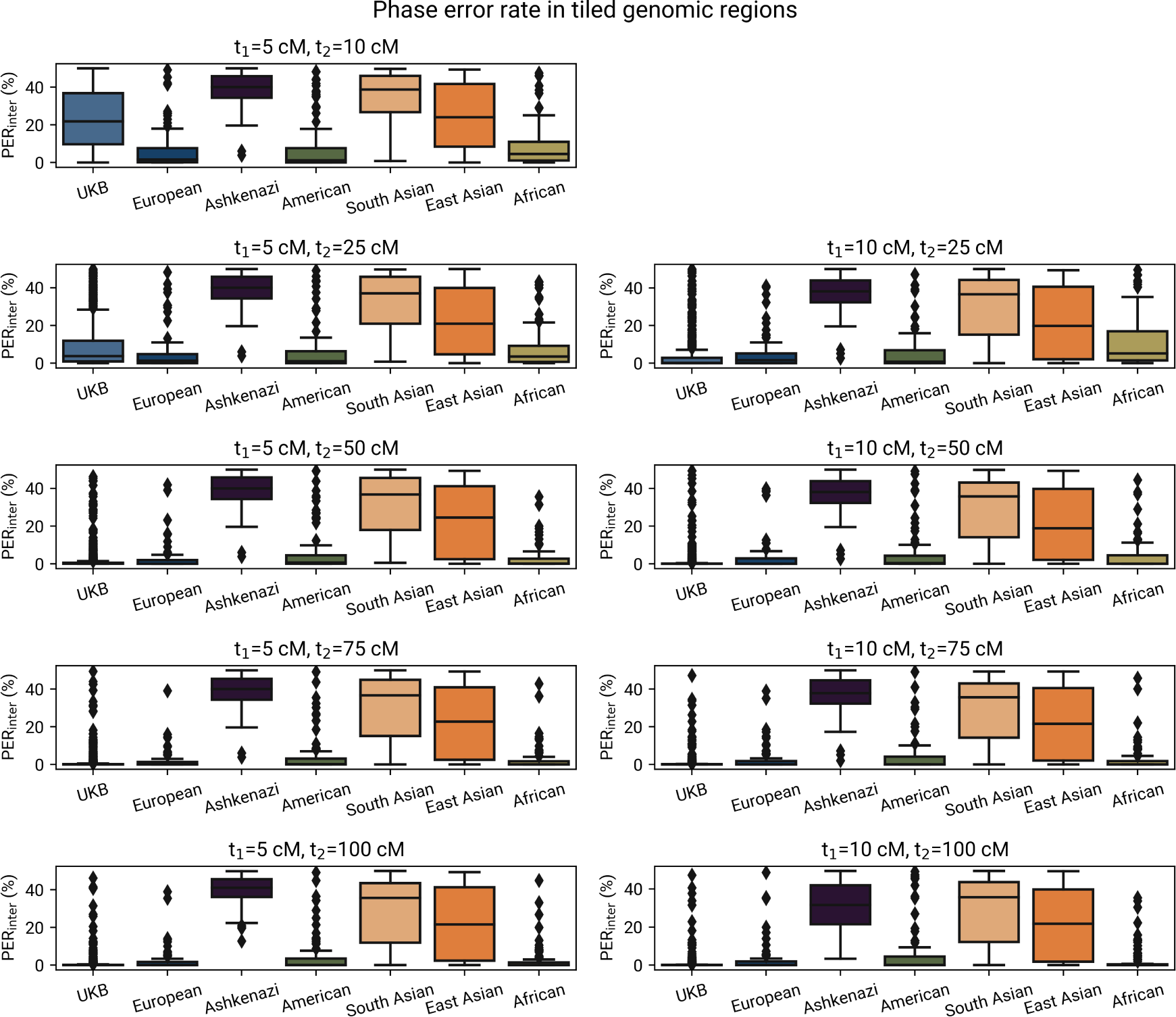
The phasing performance of HAPTIC in regions covered by IBD tiles, as measured by PER*_inter_* and split by ancestry group (all non-UKB ancestry groups are 23andMe).

**Table S1.** Switch error rate (%) of SHAPEIT and Beagle (with blips).

| Ancestry | Phaser | dataset | mean_SER | std_SER | median_SER |
| --- | --- | --- | --- | --- | --- |
| African | BEAGLE | 23andMe | 0.029019 | 0.056523 | 0.014301 |
| African | BEAGLE | UKB-TGP | 2.490410 | 1.705500 | 1.643440 |
| African | SHAPEIT | 23andMe | 0.042056 | 0.085679 | 0.025210 |
| African | SHAPEIT | UKB-TGP | 3.223650 | 1.934140 | 2.365760 |
| Ashkenazi | BEAGLE | 23andMe | 0.027088 | 0.028529 | 0.028053 |
| Ashkenazi | SHAPEIT | 23andMe | 0.043023 | 0.040968 | 0.030091 |
| East Asian | BEAGLE | 23andMe | 0.517340 | 0.612352 | 0.264857 |
| East Asian | BEAGLE | UKB-TGP | 4.921710 | 0.715112 | 5.026580 |
| East Asian | SHAPEIT | 23andMe | 0.913459 | 0.990353 | 0.521350 |
| East Asian | SHAPEIT | UKB-TGP | 5.627980 | 0.891403 | 5.728500 |
| European | BEAGLE | 23andMe | 0.033635 | 0.058727 | 0.015068 |
| European | BEAGLE | UKB-TGP | 0.709182 | 0.610696 | 0.352711 |
| European | SHAPEIT | 23andMe | 0.057813 | 0.102614 | 0.029942 |
| European | SHAPEIT | UKB-TGP | 0.978653 | 0.831223 | 0.535159 |
| admixed American | BEAGLE | 23andMe | 0.048414 | 0.117495 | 0.015017 |
| admixed American | BEAGLE | UKB-TGP | 2.041790 | 1.402580 | 1.478020 |
| admixed American | SHAPEIT | 23andMe | 0.064877 | 0.133077 | 0.028512 |
| admixed American | SHAPEIT | UKB-TGP | 2.436410 | 1.557200 | 1.839770 |
| South Asian | BEAGLE | 23andMe | 0.755690 | 0.686830 | 0.534107 |
| South Asian | BEAGLE | UKB-TGP | 2.088910 | 0.954255 | 2.190830 |
| South Asian | SHAPEIT | 23andMe | 1.216180 | 1.028100 | 0.919768 |
| South Asian | SHAPEIT | UKB-TGP | 2.758660 | 1.275120 | 3.065640 |
| UKB | BEAGLE | UKB-TGP | 0.165276 | 0.191627 | 0.116929 |
| UKB | SHAPEIT | UKB-TGP | 0.160977 | 0.198707 | 0.118048 |

**Table S2.**
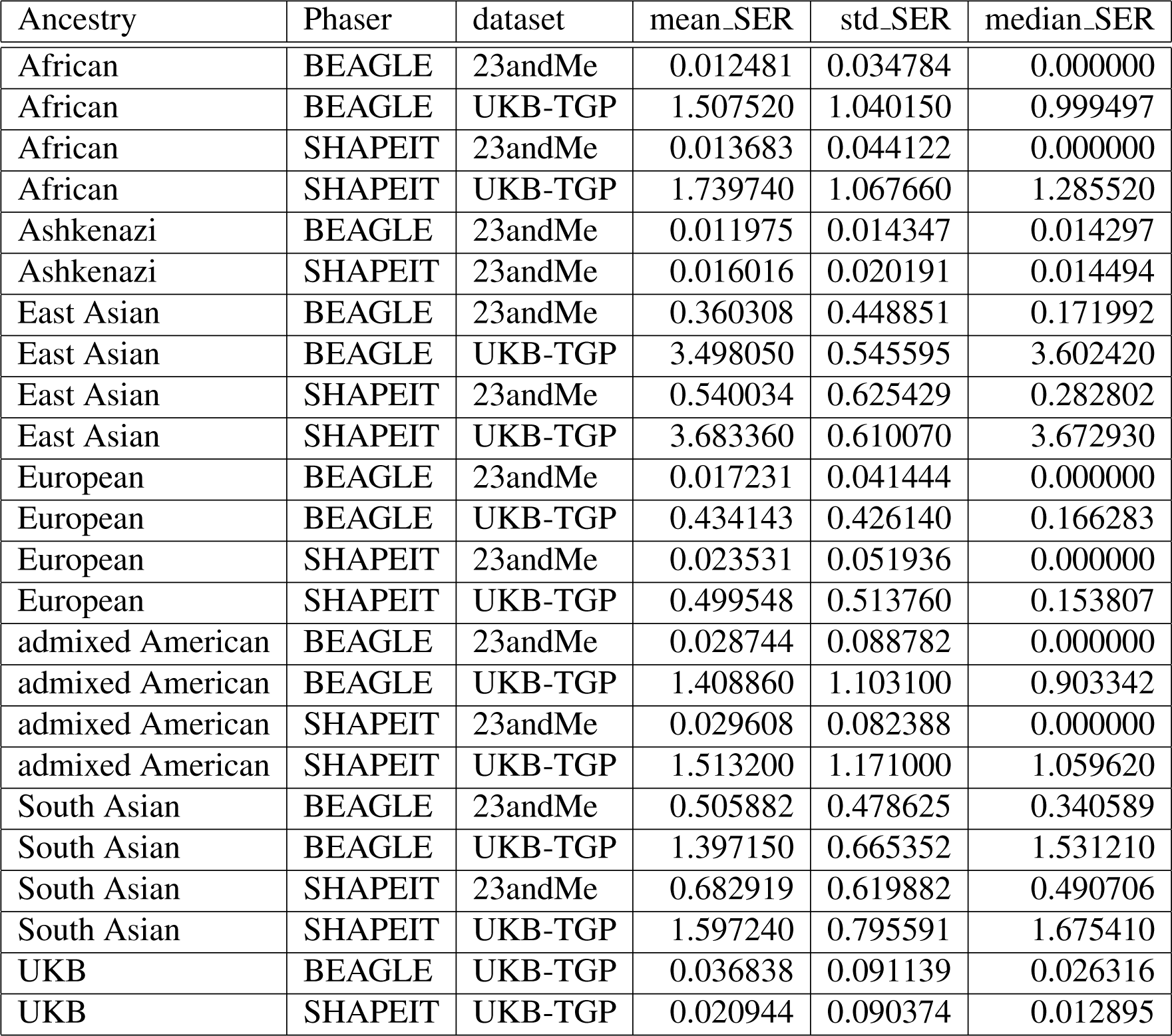
Switch error rate (%) of SHAPEIT and Beagle (excluding blips).

**Table S3.**
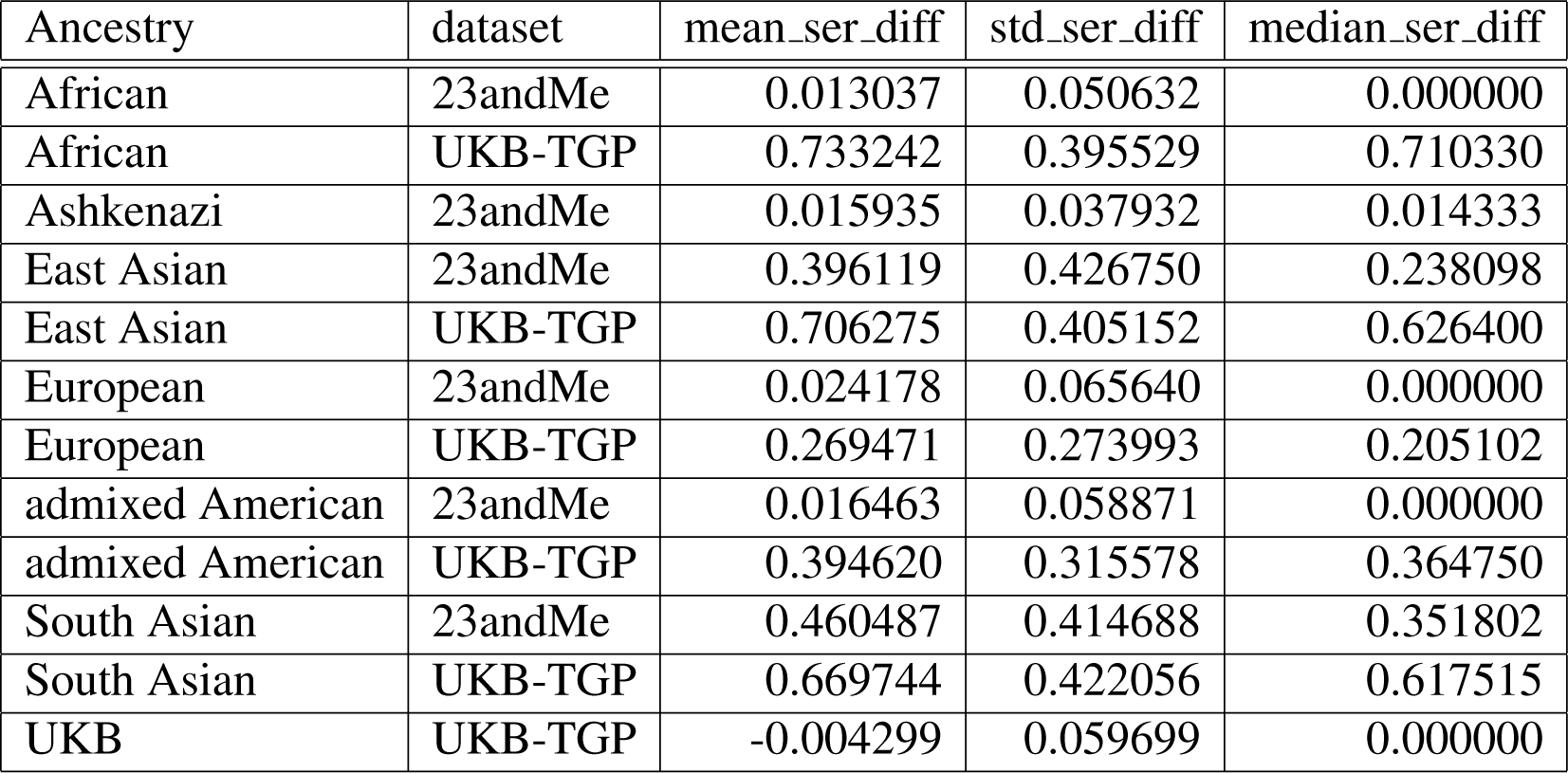
For each trio children, we looked at the difference between the SHAPEIT switch error rate (SER; %) and the Beagle SER (including blips). A positive difference indicates that SHAPEIT performed worse than Beagle. Here we show the mean, standard deviation, and median for each ancestry/dataset group.

| Ancestry | dataset | mean_ser_diff | std_ser_diff | median_ser_diff |
| --- | --- | --- | --- | --- |
| African | 23andMe | 0.013037 | 0.050632 | 0.000000 |
| African | UKB-TGP | 0.733242 | 0.395529 | 0.710330 |
| Ashkenazi | 23andMe | 0.015935 | 0.037932 | 0.014333 |
| East Asian | 23andMe | 0.396119 | 0.426750 | 0.238098 |
| East Asian | UKB-TGP | 0.706275 | 0.405152 | 0.626400 |
| European | 23andMe | 0.024178 | 0.065640 | 0.000000 |
| European | UKB-TGP | 0.269471 | 0.273993 | 0.205102 |
| admixed American | 23andMe | 0.016463 | 0.058871 | 0.000000 |
| admixed American | UKB-TGP | 0.394620 | 0.315578 | 0.364750 |
| South Asian | 23andMe | 0.460487 | 0.414688 | 0.351802 |
| South Asian | UKB-TGP | 0.669744 | 0.422056 | 0.617515 |
| UKB | UKB-TGP | -0.004299 | 0.059699 | 0.000000 |

**Table S4.**
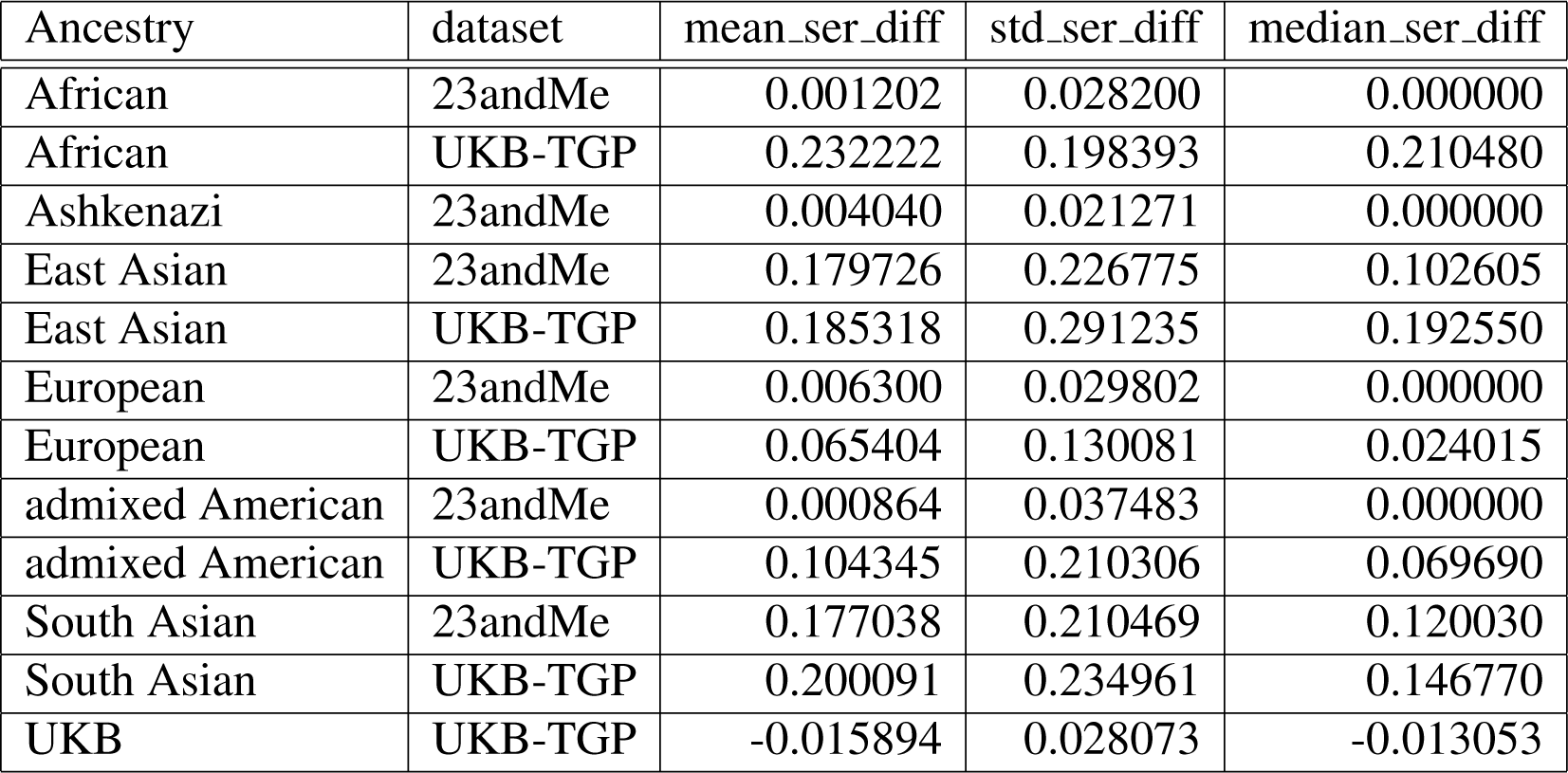
For each trio children, we looked at the difference between the SHAPEIT switch error rate (SER; %) and the Beagle SER (excluding blips). A positive difference indicates that SHAPEIT performed worse than Beagle. Here we show the mean, standard deviation, and median for each ancestry/dataset group.

**Table S5.**
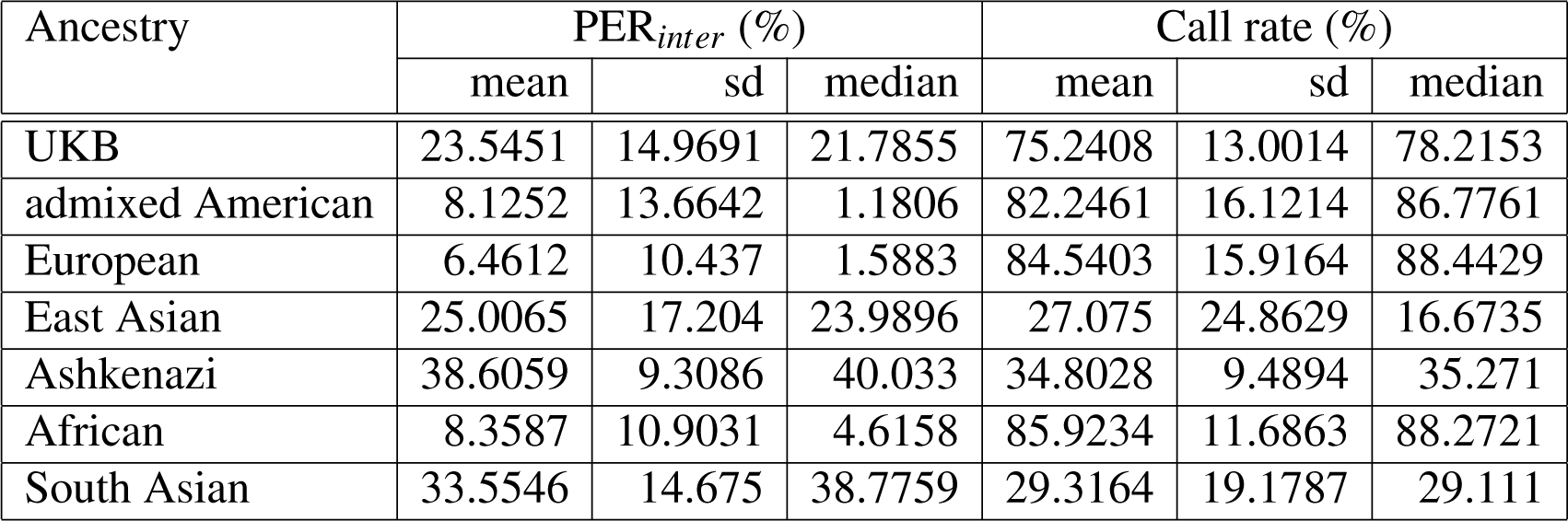
Performance of HAPTIC across different ancestry groups with *t*_1_ = 5 and *t*_2_ = 10.

**Table S6.**
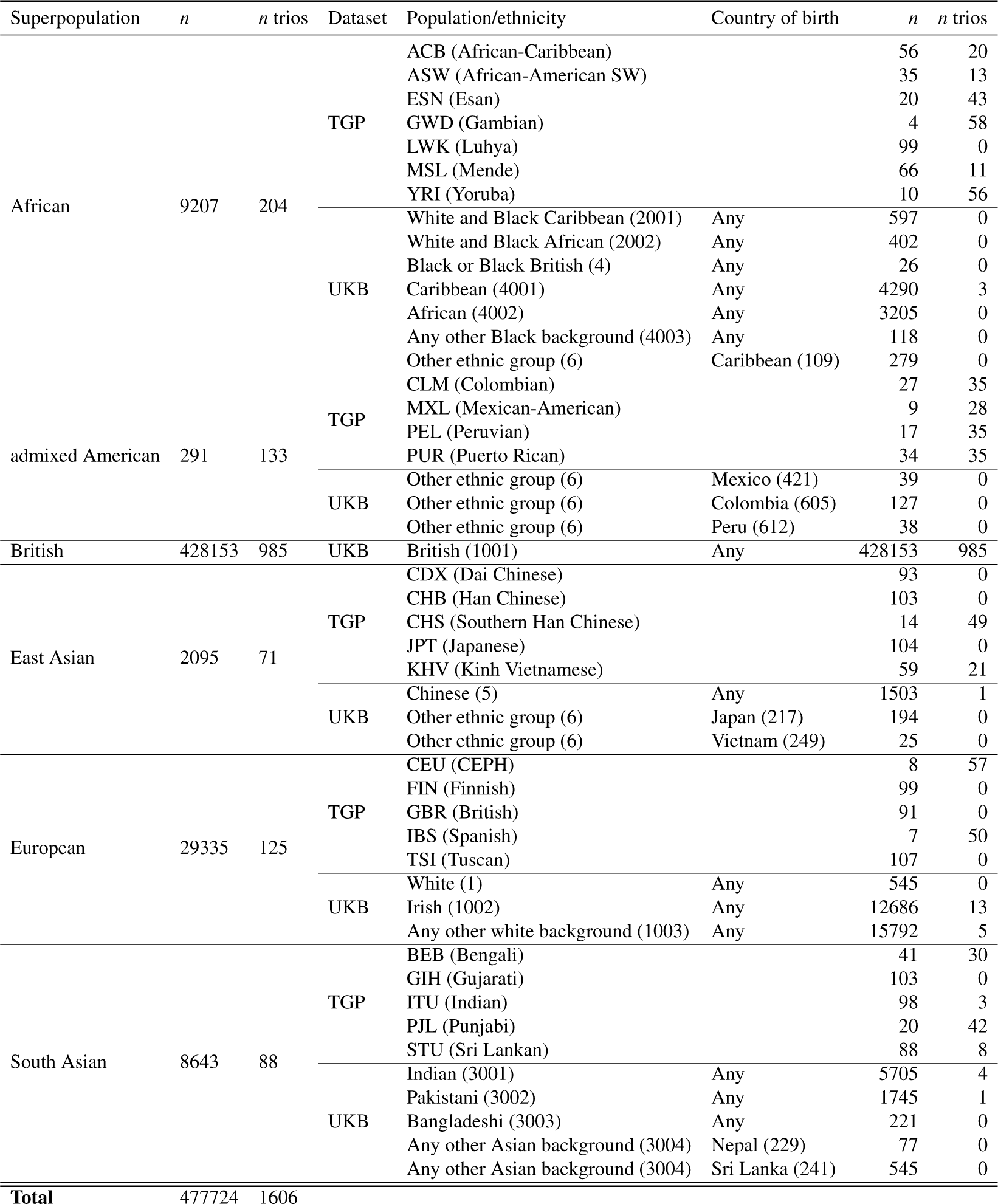
Table of the sample composition of the UKB-TGP intra-chromosomal phasing analysis. The first three columns show the Superpopulation and the aggregate counts of the total number of non-trio individuals (*n*) and the number of trios (*n* trios). The other columns show the same counts broken down by ‘sub-population’. The sub-population is defined by the ‘Population’ label of TGP samples or by the self-reported ethnicity for UKB samples if unambiguous. For a subset of UKB samples with an ambiguous self-reported ancestry (‘Other ethnic group’ or ‘Any other Asian background’), we include them if they are born in a select few countries that we attempted to match with the TGP countries (e.g., we include ‘Other ethnic group’ samples born in Mexico and classify them as American). For the UKB samples, the self-reported ethnic background is Data-Coding 1001 and the country of birth is Data-Coding 89; the codes are in the parentheses of the respective columns.

